# Asymmetric thinning of the cerebral cortex across the adult lifespan is accelerated in Alzheimer’s Disease

**DOI:** 10.1101/2020.06.18.158980

**Authors:** James M. Roe, Didac Vidal-Piñeiro, Øystein Sørensen, Andreas M. Brandmaier, Sandra Düzel, Hector A. Gonzalez, Rogier A. Kievit, Ethan Knights, Simone Kuhn, Ulman Lindenberger, Athanasia M. Mowinckel, Lars Nyberg, Denise C. Park, Sara Pudas, Melissa M. Rundle, Kristine B. Walhovd, Anders M. Fjell, René Westerhausen, the Australian Imaging Biomarkers and Lifestyle flagship study of ageing

**Affiliations:** Center for Lifespan Changes in Brain and Cognition, Department of Psychology, University of Oslo, Norway; Center for Vital Longevity, University of Texas, Dallas, TX, USA; Umeå Center for Functional Brain Imaging, and Department of Integrative Medical Biology, Umeå University, Umeå, Sweden; MRC Cognition and Brain Sciences Unit, University of Cambridge, UK; Center for Lifespan Psychology, Max Planck Institute for Human Development, Berlin, Germany; Department of Psychiatry and Psychotherapy, University Medical Center Hamburg-Eppendorf, Germany; Department of Radiology and Nuclear Medicine, Oslo University Hospital, Oslo, Norway

**Keywords:** brain asymmetry, cortical change, aging, longitudinal, lateralization

## Abstract

Normal aging and Alzheimer’s Disease (AD) are accompanied by large-scale alterations in brain organization that undermine brain function. Although hemispheric asymmetry is a global organizing feature of cortex thought to promote brain efficiency, current descriptions of cortical thinning in aging and AD have largely overlooked cortical asymmetry. Consequently, the foundational question of whether and where the cerebral hemispheres change at different rates in aging and AD remains open. First, applying vertex-wise data-driven clustering in a longitudinal discovery sample (aged 20-89; 2577 observations; 1851 longitudinal) we identified cortical regions exhibiting similar age-trajectories of asymmetry across the adult lifespan. Next, we sought replication in 4 independent longitudinal aging cohorts. We show that higher-order regions of cortex that exhibit pronounced asymmetry at age ~20 also show asymmetry change in aging. Results revealed that both leftward and rightward asymmetry is progressively lost on a similar time-scale across adult life. Hence, faster thinning of the (previously) thicker homotopic hemisphere is a feature of aging. This simple organizational principle showed high consistency across multiple aging cohorts in the *Lifebrain* consortium, and both the topological patterns and temporal dynamics of asymmetry-loss were markedly similar across replicating samples. Finally, we show that regions exhibiting gradual asymmetry-loss over healthy adult life exhibit faster asymmetry-change in AD.

Overall, our results suggest a system-wide breakdown in the adaptive asymmetric organization of cortex across adult life which is further accelerated in AD, and may implicate thickness asymmetry as a viable marker for declining hemispheric specialization in aging and AD.

**Significance:** The brain becomes progressively disorganized with age, and brain alterations accelerated in Alzheimer’s disease may occur gradually over the lifespan. Although hemispheric asymmetry aids efficient network organization, efforts to identify structural markers of age-related decline have largely overlooked cortical asymmetry. Here we show the hemisphere that is thicker when younger, thins faster. This leads to progressive system-wide loss of regional thickness asymmetry across life. In multiple aging cohorts, asymmetry-loss showed high reproducibility topologically across cortex and similar timing-of-change in aging. Asymmetry-change was further accelerated in AD. Our findings uncover a new principle of brain aging – thicker homotopic cortex thins faster – and suggest we may have unveiled a structural marker for a widely-hypothesized decline in hemispheric specialization in aging and AD.

## Introduction

Healthy aging and AD are associated with progressive disruption of structural brain organization ^1,2^ and the dedifferentiation of functionally specialized systems ^3–6^. Despite being a global organizing property of the cortex ^7,8^ with plausible relevance for hemispheric specialization ^9^, cortical thickness asymmetry has been mostly overlooked in studies examining cortical aging. Yet an asymmetrically organized cortex gives rise to efficient functional network organization ^10–13^ and thus may support cognition ^14–16^. Hence, as the cortex thins over time ^17^, cortical thickness asymmetry may also change, which may be informative for declining brain function. Here, utilizing large longitudinal datasets from 5 healthy aging cohorts and 1 dementia cohort, we aimed to establish the trajectories of asymmetric cortical thinning and examine deviations from these trajectories in AD patients.

Of the few cross-sectional studies assessing age effects upon cortical thickness asymmetry, reported results have been inconsistent ^7,14,18–20^. For example, in prefrontal regions known to be especially vulnerable in aging ^17,21^, a steeper relationship for thickness with age has been reported in both the right hemisphere (RH) ^18^ and left hemisphere (LH) ^14^ between ages 5 to 60. Still, recent large-scale meta-analyses in the ENIGMA consortium found no evidence for age effects upon frontal asymmetry ^7^, and studies have linked age to both loss ^19^ and widespread exacerbation of ^18^ thickness asymmetries established earlier in life. Conflicting results may be partly ascribable to relatively small sample sizes, and linear modelling of age effects across developmental and aging samples known to follow non-linear brain trajectories over time ^22,23^. Longitudinal lifespan studies examining intra-individual asymmetry change have the potential to resolve inconsistencies and capture non-linear asymmetry-change trajectories, but are currently absent from the literature. Thus, the foundational question of whether and where the cerebral hemispheres thin at different rates in aging remains open.

Regions of cortex that thin most in healthy aging overlap with regions exhibiting more degeneration in AD, indicating accelerated change trajectories in AD ^24–26^, and that normal and pathological brain aging may at least partly exist on a continuum ^27^. Thus, hypothesized changes in asymmetry across healthy adult life may be evident to a greater extent in AD. Although cross-sectional findings are somewhat conflicting ^19,28,29^ and meta-analyses report scant evidence for increased LH vulnerability in AD ^30^, early longitudinal findings indicate that disease progression may be associated with faster LH degeneration in medial and prefrontal cortex ^31^. Further, recent evidence hints that system-wide loss of existing asymmetries may be part of the AD phenotype ^20^. These results suggest that altered cortical asymmetry may accompany increased intra-indivdual cognitive and clinical deficit in AD over time, supporting a role for structural asymmetry in healthy brain function. Currently, however, it is not known whether purported AD-related alterations in cortical asymmetry reflect an acceleration of gradual changes occurring over the healthy adult lifespan.

Here, we tested whether the cerebral cortex thins asymmetrically in aging in an initial longitudinal adult lifespan discovery sample (LCBC; see Fig. 1A; Age-range = 20-89; total observations = 2577; longitudinal observations = 1851; mean follow-up interval = 2.7 years). To capture non-linear differences between homotopic thinning profiles across adult life, we performed vertex-wise Generalized Additive Mixed Models (GAMMs) ^32^ after resampling individual LH and RH thickness maps to a common surface (Fig.1C; SI Fig 1). Next, we applied data-driven clustering to identify regions of cortex with similar profiles of changing thickness asymmetry across adult life (Fig.1D-E). We then sought to replicate the findings in 4 independent longitudinal adult lifespan samples (see Fig.1B and SI Table 1). We hypothesized that cortical regions that are strongly asymmetric in young adulthood – as a plausible structural marker of hemispheric specialization reflecting optimal brain organization ^9^ – will exhibit changing asymmetry in aging. Finally, we predicted that similar changes in regional thickness asymmetry will occur faster in AD, and tested this using a longitudinal dataset.

**Figure 1:**
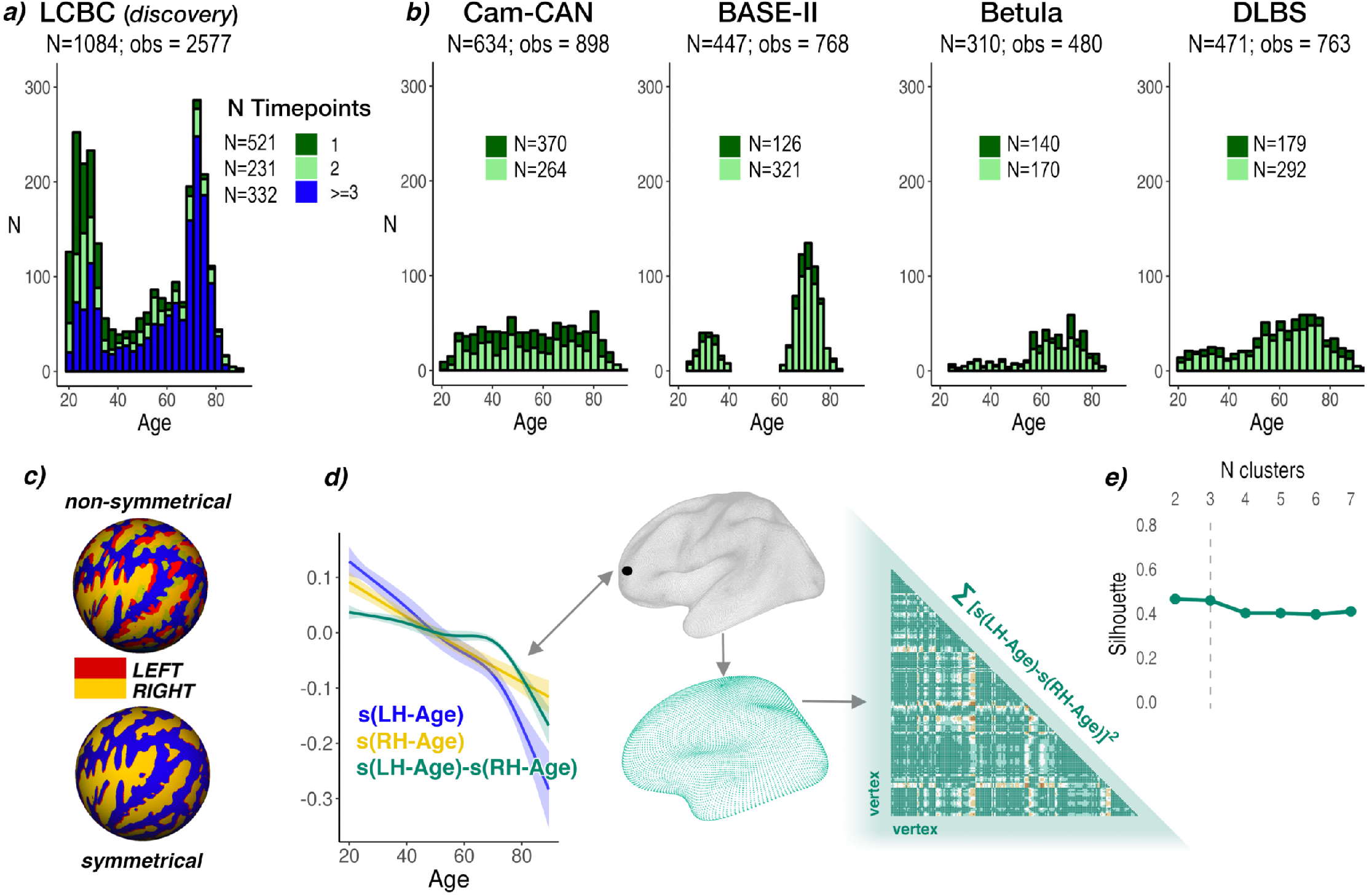
a) Age-distribution of the LCBC discovery sample. b) Age-distributions of the 4 replication samples. Colors in a) and b) b) highlight the number of longitudinal scans available per sample (i.e. dark green denotes subjects with only 1 timepoint; light green denotes subjects with 2-timepoints, blue denotes subjects with more than 2 timepoints; stacked bars). The number of unique subjects (N) are given in total and per number of available timepoints. The total number of scan observations is given per cohort (obs). c) Spherical surface rendering of a non-symmetrical (top) compared to the symmetrical template (bottom) used in the present study. Mean sulcal topography of the left and right hemispheres is shown in red and yellow, respectively. Note the near-perfect alignment of left- and right-hemisphere topography on the symmetrical surface (adapted with permission from ^33^). d) Visualization of asymmetry analysis at an example vertex (black circle). GAMMs were used to compute the zero-centered age-trajectories of the left [s(LH-Age)] and right [s(RH-Age)] hemisphere at every vertex. Blue and yellow lines indicate the smooth terms of age for the left and right hemisphere, respectively. The difference between these (green line) depicts the asymmetry trajectory [s(LH-Age)-s(RH-Age)] that was used for assessment of age-change in asymmetry. Significant asymmetry trajectories were downsampled to a lower resolution template (fsaverage5; n vertices = 10242) and a dissimilarity matrix based on the distance between trajectories at every vertex pair (computed by sum of least squares) was obtained and submitted to Partition Around Mediods (PAM) clustering. e) Mean silhouette width was used to determine the optimal partition number. Abbreviations: LH = left hemisphere; RH = right hemisphere.

## Results

### Asymmetry-change analyses

To identify cortical regions changing in thickness asymmetry across adult life, we ran vertex-wise GAMMs with Age × Hemisphere (i.e. age-related change in asymmetry) as the effect of interest in the LCBC discovery sample. Significant age-related changes in asymmetry were found in large portions of prefrontal, anterior/middle temporal, insular, and lateral parietal cortex (Fig. 2B), with particularly pronounced effects in medial prefrontal cortex (mPFC). The main effect of Hemisphere (i.e. irrespective of Age) revealed a clear anterior-posterior pattern of general thickness asymmetry in adult life (Fig. 2A). Critically, Age × Hemisphere effects conformed well to the main effect of Hemisphere, indicating that regions of cortex characterised by strong asymmetry also exhibit changing thickness asymmetry with age. Note that the direction of asymmetry-change effects is not interpretable from the GAMM interaction (see next section). An equivalent analysis with a Linear Mixed-Effect (LME) model showed comparable results (see SI).

**Figure 2:**
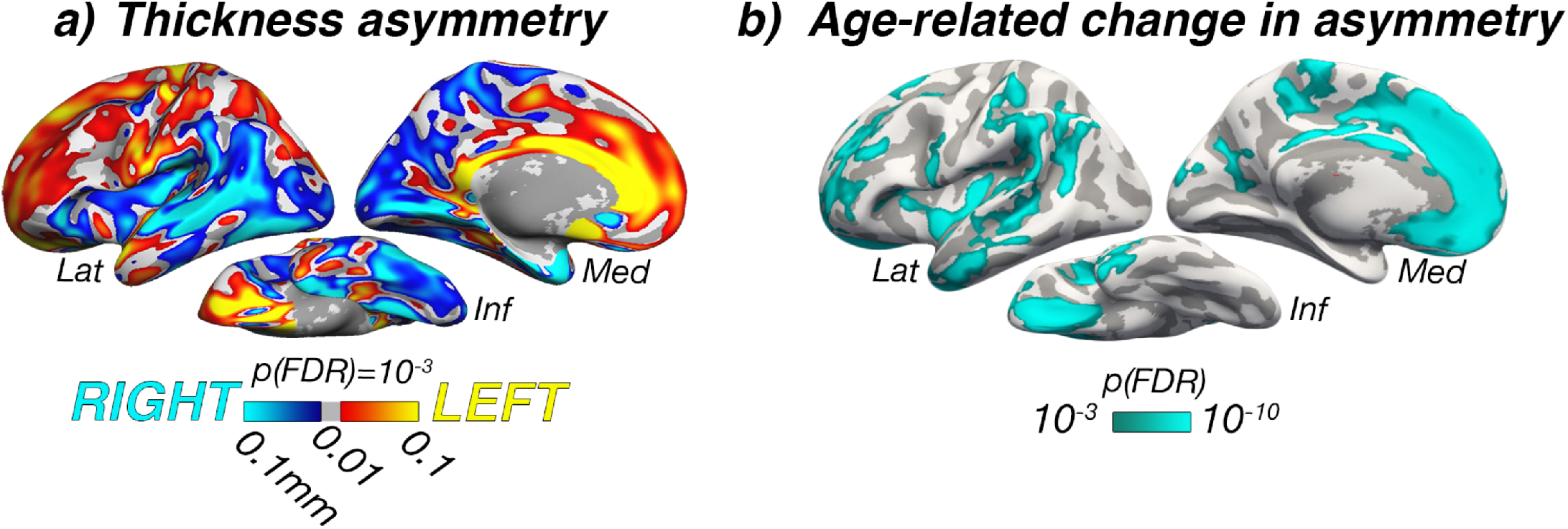
**a)** Mean thickness asymmetry for the discovery sample (2577 observations). Only regions showing significant asymmetry are shown. Warm and cold colours indicate leftward and rightward thickness asymmetry (mm), respectively. **b)** Significance map for age-changes in asymmetry. Maps are corrected for False Discovery Rate (FDR) at p < 0.001. Lat = lateral; Med = medial; Inf = inferior.

### Clustering of asymmetry trajectories

To explore asymmetry trajectories underlying the Age × Hemisphere GAMM interaction, we clustered the cortex to identify regions exhibiting similar profiles of changing thickness asymmetry across the adult lifespan. PAM clustering ^34^ indicated that asymmetry trajectories were optimally partitioned using a 3-cluster solution (mean silhouette width = .46; Fig.1E). Fig.3A-C shows the resulting 3-cluster partition. Cluster 1 consisted of vertices exhibiting mean loss of leftward asymmetry with age (negative slope from positive intercept) and mapped onto medial, orbitofrontal, dorsolateral frontal and anterior temporal cortex. Conversely, Cluster 3 vertices exhibited mean loss of rightward asymmetry with age (positive slope from negative intercept), and mapped predominantly onto insular, lateral temporoparietal cortex, and caudal anterior cingulate. Cluster 1 and Cluster 3 trajectories showed asymmetry declines emerging at a similar age (around early 30’s) and continuing across the adult lifespan, with accelerated decline around age 60 (see also SI Fig.2D). Total loss of both leftward and rightward asymmetry occurred at a similar age, around the mid 70’s. Beyond this, a continuing trend towards the opposite asymmetry to youth was apparent, particularly in Cluster 1. Cluster 2 trajectories were more mixed and generally showed weaker effects, although a trend toward loss of leftward asymmetry was still apparent. These results were robust to varying the number of clustering partitions (SI Fig.3). Homotopic age-trajectories are shown in Fig.3D for 8 main clustering-derived regions-of-interest (ROI’s) selected based on cluster- and effect-size criteria (see also SI Fig.4). Leftward asymmetry loss in frontal and anterior temporal cortex was driven by accelerated LH-thinning. Conversely, rightward asymmetry loss was driven by accelerated RH-thinning (e.g. in insular and lateral parietal cortex; Fig.3D). Only the cingulate region did not conform to this pattern, exhibiting an early leftward asymmetry that increased with age. Importantly, accelerated thinning corresponded to the percentage of cortex lost with age (i.e. hemispheric differences in rates-of-thinning were relative; also indexed by crossing trajectories). Overall, findings in the discovery sample indicate that changes in cortical thickness asymmetry are a feature of aging, and that accelerated thinning of the thicker hemisphere leads to extensive loss (or reversal) of asymmetry in older age.

**Figure 3:**
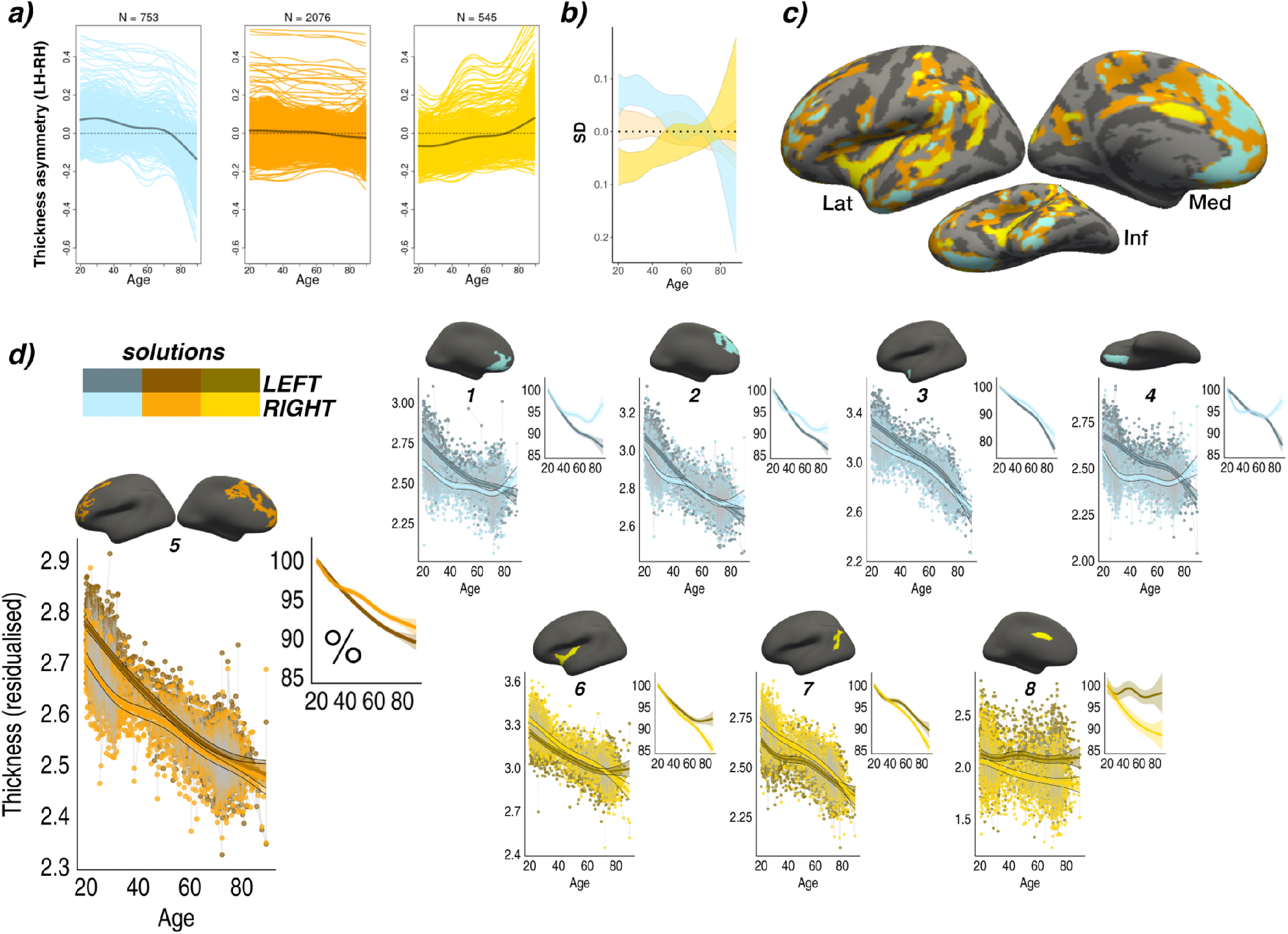
**a)** PAM clustering solutions for asymmetry trajectories in the LCBC discovery sample. Each colored line represents a vertex. Mean trajectories are in grey. Vertices showing leftward asymmetry in early adult life (higher than dotted line) typically exhibit loss of leftward asymmetry with age (blue plot), whereas vertices showing rightward asymmetry (lower than dotted line) typically exhibit loss of rightward asymmetry (yellow plot). Importantly, because asymmetry trajectories were computed as the difference between zero-centered hemispheric trajectories [s(LH-Age)-s(RH-Age)] (cf. Fig 1D), mean differences between LH and RH thickness (i.e. mean asymmetry/the intercept) are not taken into account and do not influence the clustering. For visualization, we added the main effect of Hemisphere, vertex-wise, and computed the mean asymmetry trajectory of vertices in each clustering solution. **b)** Standard deviations (SD) of the asymmetry trajectories for the clustering solutions. **c)** Solutions mapped on the surface. **d)** Thinning trajectories plotted separately for LH and RH in regions derived from the clustering (numbered 1-8). Colors correspond to the solutions in a), and darker shades indicate LH trajectories. All trajectories were fitted using GAMMs. Data is residualized for sex and scanner. Except where otherwise stated, ribbons depict 95% confidence intervals. Smaller plots illustrate percentage-change with age for each region.

### Replication analyses

We next tested whether the clustering of asymmetry trajectories was reliable across 4 independent longitudinal adult lifespan cohorts (Cam-CAN, BASE-II, BETULA and DLBS: described in SI Table 1). Clustering solutions in Cam-CAN, BASE-II and BETULA exhibited highly similar mean intercept and slope effects to those described in the discovery sample, and mapped onto near-identical regions (Fig. 4). In these, asymmetry-loss effects conformed to the anterior-posterior pattern of general cortical asymmetry in the same manner as in the discovery sample. Quantitatively, the clustering trajectories in Cam-CAN, BASE-II and BETULA were comparable to the clustering trajectories in LCBC (SI Fig. 5-6). Spatially, the similarity of clustering results to those observed in the discovery sample was substantiated by Dice coefficients far higher than expected by chance (Cam-CAN = .54; BASE-II = .56; BETULA = .50; DLBS = .41; true expected Dice at random = .1, all *p_perm_* < .001). The clustering in DLBS showed only partial replication: contrary to other samples, mPFC and temporal cortex clustered with regions characterised by mean rightward asymmetry loss, and intercept effects indicative of early leftward asymmetry were not evident in the estimated asymmetry trajectories in DLBS. Vertex-wise GAMM effects also appeared largely consistent across samples (see SI). Overall, the clustering of thickness asymmetry age-trajectories in the LCBC discovery sample replicated in 3 of 4 longitudinal aging cohorts (see SI Figs 8-9 for full results varying the number of clustering partitions in each cohort).

**Figure 4:**
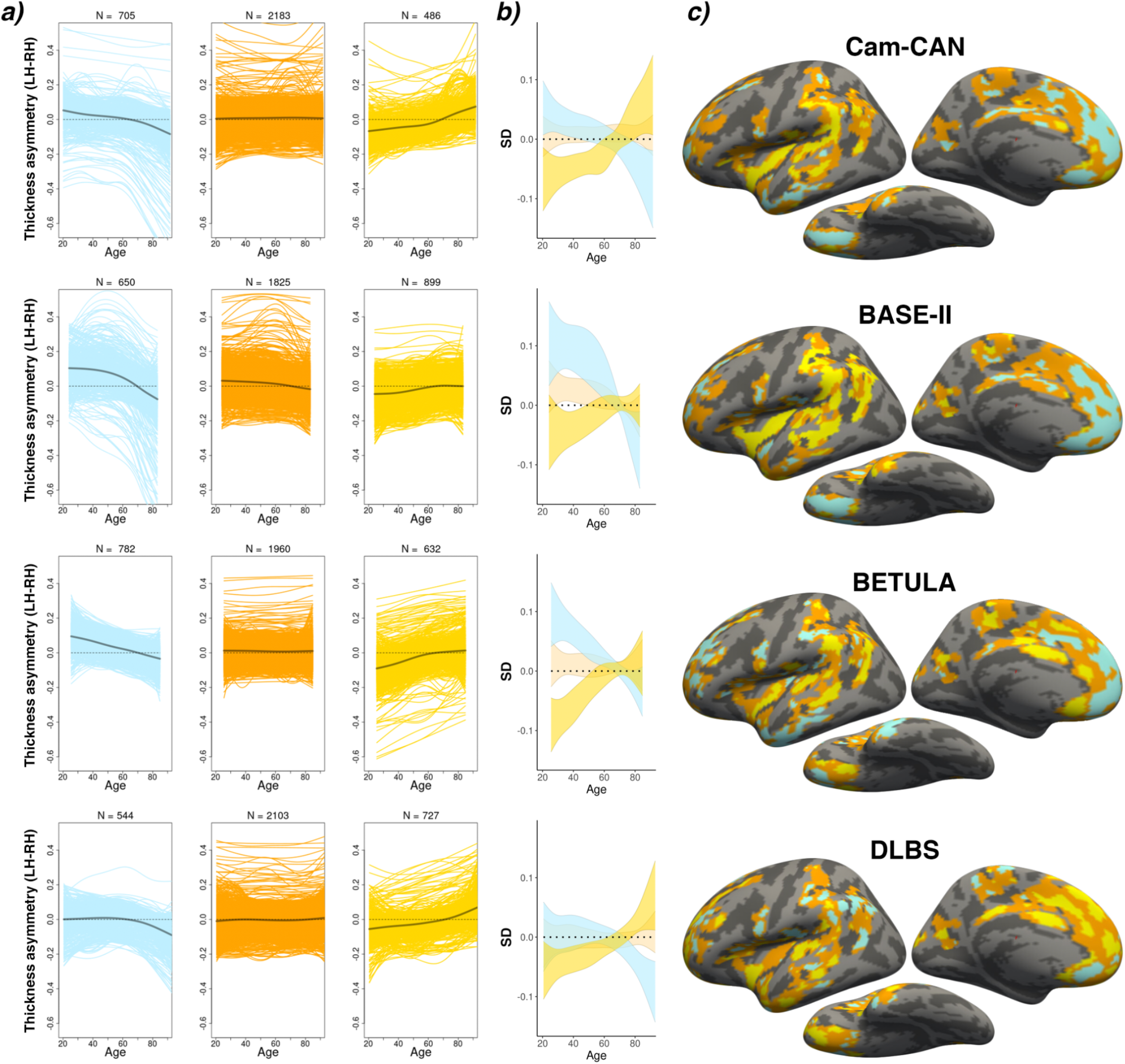
**a)** PAM clustering solutions of asymmetry trajectories in the four replication samples. Each colored line represents a vertex. Mean trajectories are in grey. In three out of four samples, the clustering solutions were highly similar to the discovery sample (cf Fig. 3; SI Fig. 5). **b)** Standard deviations (SD) of the asymmetry trajectories for the clustering solutions. **c)** Solutions mapped on the surface. Results are depicted row-wise per cohort. All trajectories were fitted using GAMMs.

### Cognitive change analysis

We next a ran GAMM for each ROI to assess whether longitudinal thickness asymmetry change predicted longitudinal cognitive change in verbal memory and fluid reasoning ability – cognitive domains that show high vulnerability in aging ^35^. While age explained 29% and 36% of the variance in longitudinal memory and fluid reasoning scores, respectively (sex controlled; SI Fig.10; SI Table 3), we observed no significant (pFDR < .05) effects of thickness asymmetry change on longitudinal change for either cognitive measure in any of the 8 ROIs (SI Table 4). Further, including all 8 asymmetry ROI’s in a GAMM did not significantly improve model fit, suggesting that asymmetry-change had no additive effect upon longitudinal cognitive change (verbal memory *p* = .98; fluid reasoning *p* = .79; see SI).

### Longitudinal AD analysis

Finally, we asked whether in AD, asymmetry-loss is accelerated in regions prone to exhibit asymmetry-change in healthy aging. We used data from the Australian Imaging Biomarkers and Lifestyle study of aging (AIBL; longitudinal observations only) to define longitudinal groups of healthy aging and AD indivduals (see Fig.6A and Methods), and ran LME analyses for each of the 8 ROI’s testing for group differences in asymmetry-change over time. AD individuals were found to exhibit a steeper decline of (leftward) thickness asymmetry over time compared to cognitively healthy controls in most frontal cortical regions, and in anterior temporal cortex (see SI Table 5 for full results). Whether or not an ROI showed opposite asymmetry in AD at baseline (i.e. thicker RH due to LH depletion) corresponded with whether the ROI lifespan trajectories showed opposite asymmetry in older age (cf Fig. 3D; superior mPFC and orbitofrontal ROI’s). Overall, the analyses confirmed that frontal and temporal patterns of asymmetry-change across healthy adult life are accelerated in AD.

**Fig.6.**
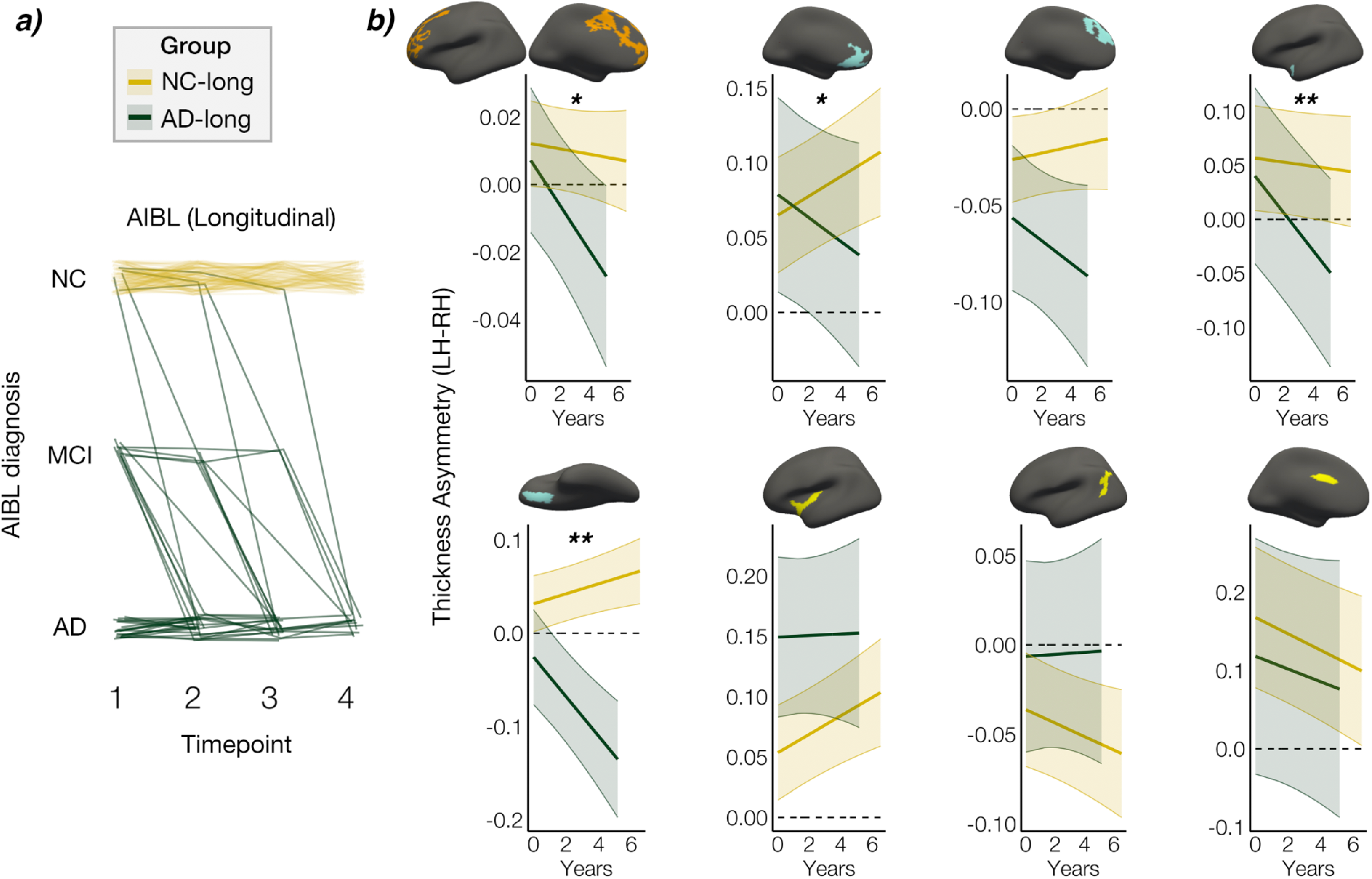
Thickness asymmetry change in AD versus healthy aging. A) X-axis denotes study timepoints. Single-timepoint diagnoses (y-axis; NC = normal controls; MCI = Mild Cognitive Impairment; AD = Alzheimer’s Disease) were used to define two longitudinal Groups of AD and NC individuals. Each line represents a subject and the color denotes longitudinal group membership (AD-long, in green; NC-long, in gold). AD-long individuals were diagnosed with AD by their final timepoint whereas NC-long were classified as healthy at every timepoint. Note that single-timepoint MCI diagnoses were consisdered only for the purpose of defining the longitudinal AD group (see methods). B) Interaction between Group and Time (years) since baseline measurement. Bands represent 95% confidence intervals. Accelerated change in thickness asymmetry was observed in the AD group in frontal and anterior temporal cortical ROI’s. p(FDR) * <.05; ** <.01.

## Discussion

We discovered that the cerebral cortex thins asymmetrically across adult life. Age-changes in asymmetry almost invariably reflected progressive asymmetry-loss and hence were precipitated by faster thinning of the (previously) thicker homotopic hemisphere. In four out of five longitudinal adult lifespan samples the cortex clustered into a highly reproducible pattern that described age-trajectories of asymmetry topologically across the cortex. This mapped onto a general anterior-posterior pattern of left-right thickness asymmetry and revealed loss of both leftward and rightward asymmetry on a similar time-scale across adult life, suggesting system-wide loss of asymmetry in aging. Finally, we found that frontal and temporal regions vulnerable to asymmetry-loss in healthy aging exhibit accelerated asymmetry change in AD.

As we demonstrated, regional changes in cortical thickness asymmetry are an overlooked yet important feature of normal brain aging that is both shared by and accelerated in neurodegenerative AD. Across multiple cohorts, loss of leftward asymmetry over healthy adult life was particularly pronounced in frontal and anterior temporal cortex, and loss of rightward asymmetry was evident in insular and lateral temporoparietal regions. The average temporal dynamics of asymmetry-loss were also markedly preserved across samples, with gradual asymmetry decline in early adulthood, and accelerated decline from age ~60 onwards. These trajectories applied to asymmetry fit well with general thinning trajectories known to show accelerated decline from age 60 onwards ^36,37^, though we note that individual regional trajectories did not always conform to the average time-course. Regions we observed changing in asymmetry were predominantly located in higher-order association cortex, which has been found to exhibit less average interhemispheric connectivity ^38^ in favour of more lateralized within-hemisphere interactions ^10^ – supporting notions of hemispheric specialization ^9,12^. Speculatively, our observation that thicker homotopic cortex thins faster may be in line with contemporary evidence indicating that age pathologies propogate transneuronally though synaptic connections ^39,40^ which may precipitate faster degeneration of the hemisphere with more cortico-cortical connections ^41^. As evidence suggests cortical thinning partly reflects synaptic loss ^42–44^, critical questions for future research concern the underlying neurobiological mechanisms, exact functional consequences, and indeed whether the principle that thicker cortex thins faster only applies to homotopy. And while it remains unclear how the phenomenon may be reflected in age-related asymmetry differences in electrophysiological ^45^ and functional magnetic resonance signals ^46,47^, our findings may be in line with studies indicating decreased left-lateralization of frontal resting-state networks with higher age ^48^. Regardless, our results illustrate a hitherto undescribed phenomenon that fits within the well-established dedifferentiation view of the aging brain ^3–5^: the mirrored reduction of leftward and rightward asymmetry across adult life suggests a system-wide breakdown in the adaptive asymmetric organization of the cortex in aging.

An asymmetrically organized cortex leads to increased proximity of collaborating brain regions and intrahemispheric clustering of specialized networks ^11–13^. The asymmetry-loss observed here may reflect decline in the intrahemispheric network efficiency of higher-order regions which are more inhibited from cross-hemispheric interaction ^10,38^ via the corpus callosum ^49,50^. Because higher-order regions may also be more prone to homotopic disconnection with age ^51^, future work assessing how age-changes in thickness asymmetry relate to altered interhemispheric connectivity and callosal degeneration ^52–54^ may shed light on important structure-function relationships in brain asymmetry and aging. Recent large-scale meta-analyses found evidence for age effects on thickness asymmetry only in superior temporal cortex ^7^ and previous cross-sectional studies have yielded mixed results ^14,18,19^. Thus, by highlighting asymmetry-loss as a system-wide process in aging we here substantially extend previous knowledge. This also highlights the advantage longitudinal aging studies hold over meta-analyses based on cross-sectional age models in samples of varying size ^55,56^, as well as the advantage of vertex-wise asymmetry approaches. The results presented here fit with the view that brain systems subserving higher-level associative cognition in particular become less specialized and disorganized in aging ^57^.

The reliable anterior-posterior pattern of general thickness asymmetry found here (see SI Fig. 7) agrees with previous work ^8,14^, has recently been shown in global meta-analyses ^7^, and is compatible with reports of developing thickness asymmetry from birth ^58,59^. Reproducibility across samples and emerging cross-study consensus also in early development suggests a genetic influence upon cortical thickness asymmetry ^58^, and genetic factors have recently been implicated in the dynamics of age-related cortical change across life ^36^. One can speculate whether age-related asymmetry breakdown is a by-product of genetic factors encoding the asymmetric organization of cortex ^60^ supporting hemispheric specialization of function, and that this differentiation leads to downstream neural consequences in aging. Though anatomo-functional relationships are likely complex ^61^, our results suggest that cortical thickness asymmetry may constitute a viable anatomical marker for key aspects of human hemispheric specialization. For example, we found robust evidence that mPFC asymmetry is particularly vulnerable in aging. Although implicated in a wide variety of complex cognition, the role of mPFC in executive function and normal memory operations is well-established ^62,63^, and deficits in these are considered hallmarks of aging ^35^. Although future research is needed to assess the specific cognitive relevance of regional thickness asymmetries and structure-function change relationships, the profound asymmetry-loss observed here raises the possibility that complex cognitive abilities susceptible to decline in most individuals with advancing age may at least be partly subserved by hemispherically specialized networks.

Indeed, we found that asymmetry-change extended through the aging-neurodegeneration continuum. Specifically, we observed accelerated loss of LH frontal and temporal cortices above and beyond those observed during healthy aging, illustrating that gradual age-related changes in asymmetry are exacerbated in AD. This agrees with previous longitudinal evidence indicating faster LH neurodegeneration in AD^31^, recent work hinting at systemic AD-related asymmetry-loss ^20^, and the regional susceptibility of frontal and temporal cortices to AD pathology ^24,25^. One can speculate whether and to what degree faster LH degradation tracks to an asymmetric presence of other AD biomarkers, such as neurofibrillary tangles ^64^, as patterns of cortical thinning in AD largely overlap with tau deposition ^25,65^. Future research could also assess whether and how cortical asymmetry change in AD might co-occur with asymmetric neurodegeneration of subcortical brain structures vulnerable in AD ^66,67^.

Here, we show the notion that characteristic changes in cortical structure in AD are also observed, though to a minor degree, in cognitively normal aging can also be extended to include thickness asymmetry ^24–26^. Hence, the present findings suggest a continuity of change between healthy and pathological brain aging, and highlight the importance of lifespan perspectives for understanding the pathophysiology of AD.

The implication that thickness asymmetry is important for healthy brain function agrees with a recent meta-analysis confirming that subtly reduced asymmetry is a feature of neurodevelopmental disorders ^16^. However, our results also indicate that individual differences in regional asymmetry-change had little predictive value upon longitudinal cognitive scores across cognitively healthy adult life (neither memory nor fluid reasoning ability). Thus, loss of thickness asymmetry may be more relevant for advanced cognitive decline – such as that associated with AD – but less sensitive to individual differences in cognitive decline trajectories across the healthy adult lifespan. Although we note that alternative tests such as cognitive speed measures may be better suited than task accuracy for assessing asymmetry-cognition relationships, our results suggest that asymmetry-change may be unlikely to be a sensitive marker for age-related decline at the individual level (similar conclusions have recently been drawn in relation to asymmetry in other contexts ^68^). This is also evidenced by the fact that our large longitudinal sample sizes were needed to expose small-to-medium effects that nevertheless translated to consistent gradual changes in asymmetry across adult lifespan samples.

Some potential caveats should be considered. First, the estimation of GAMM age-trajectories will be affected by age-distribution because knot placement is based upon data density, which differed across discovery and replication samples. This may explain some heterogeneity in vertex-wise results between samples (SI Fig.6) and in the subsequent clustering of asymmetry trajectories, and also limits the interpretation and replication of exact timings (e.g. acceleration and inflection points) of asymmetry change across adult life. Relatedly, survivor bias in longitudinal aging studies will also affect the estimation of lifespan trajectories, and this could be one candidate explanation for the lack of full replication in DLBS. That is, if DLBS is more biased to recruiting high performing older adults, this could preclude replication of the lifespan trajectories, and varying degrees of age-related pathology in later life could explain some of the differences in the lifespan reduction of asymmetry observed across samples. This would also agree with our observation that individuals who either had or were later diagnosed with AD (and hence were classed as cognitively normal at one or more timepoints) exhibited a higher reversal of the group-average asymmetry pattern at baseline. Second, because GAMMs estimate trajectories as a combination of longitudinal and cross-sectional effects, the inclusion of more time-points and longer follow-up intervals will better approximate the true longitudinal trajectories. Third, to ensure clustering reliability we excluded small clusters assumed to be more prone to noise. Yet we also observed consistent vertex-wise effects in smaller regions of cortex that violated this assumption (e.g. parahippocampal gyrus; SI Fig.6), and thus cannot exclude other, more focal changes in asymmetry potentially informative for cognitive decline in aging. Third, because the clustering distributes all vertices among a given number of solutions, the trajectories of some vertices may not fit well within a given solution despite statistically fitting best to that solution. Consequently, the cluster solutions largely inform about the average trends rather than the asymmetry trajectories of individual regions, which showed more heterogeneity in shape. This forcing may also explain some spatial heterogeneity in clustering across samples. Nevertheless, the clustering protocol – which was based solely on the age-trajectories of asymmetry and was blind to the spatial location in cortex – allowed for a concise description and comparison of the average asymmetry trajectories evident in each aging cohort.

## Conclusion

Brain asymmetry seems to have arisen under evolutionary pressure to optimize processing efficiency and is broadly thought to confer organizational advantages that benefit brain function. Here, we show that the asymmetric organization of higher-order cortical regions in young adulthood decays with advancing age, and this decay follows a simple general organizing principle: the thicker of the two homotopic cortices thins faster. This principle was highly reproducible across different cohorts and was significantly accentuated in AD patients. Overall, the present study may have unveiled the structural basis of a widely suggested system-wide decline in hemispheric functional specialization across the adult lifespan, and found a potential continuation and acceleration of this decline in AD.

## Method

### Samples

#### Discovery sample

An adult lifespan sample comprising 2577 scans from 1084 healthy individuals aged 20.0 to 89.4 (703 females) was drawn from the Center for Lifespan Changes in Brain and Cognition database (LCBC; Department of Psychology, University of Oslo). See SI for details. All individuals were screened via health and neuropsychological assessments. The following exclusion criteria were applied across LCBC studies: evidence of neurologic or psychiatric disorders, use of medication known to affect the central nervous system (CNS), history of disease/injury affecting CNS function, and MRI contraindications. Participants in LCBC studies were required to consider themselves right-handed. Additionally, the following inclusion criteria were defined here: < 21 on the Beck Depression Inventory ^69^, and ≥ 25 on the Mini Mental Status Exam ^70^.

#### Replication samples

Four longitudinal adult lifespan datasets were selected to test reliability of the discovery sample findings. Three were derived from the *Lifebrain* consortium – a data-sharing initiative between major European lifespan cohorts ^56^: the Cambridge Center for Aging and Neuroscience study (*Cam-CAN;* N = 634; N scans = 898) ^71,72^, the Berlin Study of Aging-II (*BASE-II;* N = 447; N scans = 768) ^73^ and the *BETULA* (N = 310; N scans = 480) project ^74^. The fourth was the Dallas Lifespan Brain Study (*DLBS;* N = 471; N scans = 763) ^75^. Longitudinal data (up to two-timepoints) was sourced for each replication cohort (see SI Tables 1-2 for sample demographics and MRI image parameters).

#### AD sample

Data was collected by the AIBL study group (see ^76^). We used the single-timepoint AIBL diagnosis (normal controls [NC]; Mild Cognitive Impairment [MCI]; Alzheimer’s Disease [AD]) to identify 2 longitudinal groups based on final-timepoint diagnosis: *NC-long* consisted of subjects classed as NC at every timepoint; *AD-long* consisted of subjects classed as AD throughout or by the final timepoint (see Fig. 6A). Individuals reverting from MCI or AD diagnosis were not included. 545 observations of 169 subjects were included in the sample (timepoint range = 2-4; see SI Tables 1-2).

### MRI acquisition

Discovery sample (LCBC) data consisted of T1-weighted (T1w) magnetization prepared rapid gradient echo (MPRAGE) sequences collected on two scanners at Oslo University Hospital; a 1.5 Tesla (T) Avanto and a 3T Skyra (Siemens Medical Solutions, Germany). The number of Avanto and Skyra T1w scans was 832 and 1745, respectively. MRI acquisition parameters for all samples are summarized in SI Table 3.

### MRI preprocessing

Cortical reconstruction of the anatomical T1-weighted images was performed with FreeSurfer’s longitudinal pipeline (v6.0.0, http://surfer.nmr.mgh.harvard.edu/wiki) ^77^ on the Colossus computing cluster at the University of Oslo. This fully automated pipeline yields a reconstructed surface map of cortical thickness estimates for each participant and timepoint using robust inverse consistent registration to an unbiased within-subject template ^78^ (see SI). A high-resolution symmetrical surface template (“*LH_sym*”)^33^ was used to resample the FreeSurfer-estimated cortical thickness maps of the LH and RH of each participant into a common analysis space. This procedure achieves precise vertex-wise alignment between cortical hemispheres (see Fig.1C; SI Fig.1). *LH_sym* was created from a composite of LH and RH surface models in a database enriched in left-handers: the BIL&GIN ^33,79^. In symmetrical space, an 8 mm full-width-half-maximum Gaussian smooth-kernel was applied across the surface to the LH and RH thickness data.

### Asymmetry-change analyses

The analysis pipeline is summarized in Fig. 1D. Thickness maps were analyzed vertex-wise using GAMMs implemented in R (v3.5.0; “gamm4” package; ^80^). We used a factor-smooth GAMM interaction approach that allowed fitting a smooth Age-trajectory per Hemisphere and testing the smooth Age × Hemisphere interaction. Hemisphere was additionally included as a fixed-effect, Sex and Scanner as covariates-of-no-interest, and a random subject intercept was included. To minimize overfitting the number of knots was constrained to be low (k=6). Significance of the smooth Age × Hemisphere interaction was formally assessed by testing for the existence of a difference between the smooth term of Age across Hemispheres (see SI code). FDR-correction controlling for positive dependency ^81^ was applied to the resulting Age × Hemisphere and Hemisphere (main effect) significance maps, and the maps were masked at p(FDR) < 0.001.

### Clustering of asymmetry age-trajectories

The linear predictor matrix of the GAMM was used to obtain *asymmetry trajectories* [*s(LH-Age)-s(RH-Age)*] underlying the interaction, computed as the difference between zero-centered hemispheric trajectories. After removing the smallest clusters (<300mm^2^), we resampled significant asymmetry trajectories to a lower resolution template (*fsaverage5*), calculated a dissimilarity matrix (see Fig. 1D) and submitted it to PAM clustering (“cluster” R package; ^34^). Clustering solutions from 2 to 7 were considered, and the optimal solution was determined by the mean silhouette width. The largest cortical regions within each clustering solution were considered as ROI’s in post-hoc analyses (13 initial ROIs). Individual-level LH and RH thickness maps were then resampled from the symmetrical space to *fsaverage5,* and mean thickness was extracted for each ROI. To describe the patterns underlying the smooth interaction, for each ROI we ran comparable factor-smooth GAMMs and computed LH and RH trajectories. We then re-ran these after removing outliers (both hemispheres; defined as residuals ≥ 6 SD from the fitted trajectory of either hemisphere), and plotted the fitted Age trajectories for each hemisphere. Relative change was calculated by scaling fitted trajectories by the prediction at the minimum age. 8 main ROI’s were identified to be used in subsequent ROI-based analyses based upon size and degree of asymmetry (see SI Fig.2).

### Replication analysis

The GAMM analysis and clustering protocol was repeated for each longitudional aging cohort. To evaluate clustering similarity the clustering was constrained within the same set of vertices as the discovery sample. For each clustering solution, Dice similarity was computed between discovery and replication maps, and the mean Dice across all solutions was taken. Significance of the spatial overlap in clustering results between discovery and replication samples was assessed across 10,000 random permutations of the clustering solution maps. Equivalent vertex-wise factor-smooth GAMM analyses were ran for each sample independently - minus the Scanner covariate (all were single-site).

### Cognitive-change analysis

We considered longitudinal scores on the California Verbal Learning Test (CVLT) ^82^ and the Matrix Reasoning subtest of the Weschler Abbreviated Intelligence Scale (WAIS) ^83^ as proxies of memory- and fluid reasoning ability-change, respectively. We used a subset of the LCBC sample for which at least two timepoints (range = 2-4) of cognitive tests were available. The total number of observations for CVLT and matrices with concurrent scans were 783 (N = 312) and 788 (N = 314), respectively (mean N timepoints = 2.7; mean follow-up interval = 3.2 years; max interval = 10.9 years). For matrices we used the raw scores. For CVLT, we used the first principal component across z-transformed raw scores on the learning, immediate, and delayed free-recall subtests. Next, for each of the 8 main ROI’s we ran GAMMs predicting each cognitive measure with the following smooth terms: Asymmetry (LH-RH), mean thickness (across hemispheres) and Age (Sex, Scanner, Test Version controlled; random subject intercept).

### Longitudinal AD analysis

For AIBL data (28^th^ April 2015 release) we calculated thickness asymmetry (LH-RH) on *LH_Sym*, downsampled to *fsaverage5*, and extracted from clustering-derived ROIs. For each ROI, we ran LME’s with the factors Group and Time (from baseline), and assessed the Group × Time interaction to test for group differences in asymmetry-change. Age (at baseline), Sex and Scanner were included as covariates, and a random subject intercept was included. As only year-of-birth data is available with AIBL, Age was estimated by randomly jittering the interval between the halfway date in the year-of-birth and test date.

## Funding

The Lifebrain project is funded by the EU Horizon 2020 Grant: ‘Healthy minds 0–100 years: Optimising the use of European brain imaging cohorts (‘Lifebrain’).” Grant agreement number: 732592. In addition, the different sub-studies are supported by different sources: LCBC: The European Research Council under grant agreements 283634, 725025 (to A.M.F.) and 313440 (to K.B.W.), and the Norwegian Research Council (to A.M.F., K.B.W.) under grant agreement 249931 (“TOPPFORSK”), The National Association for Public Health’s dementia research program, Norway (to A.M.F).

## Acknowledgements

The authors are indebted to Fabrice Crivello, Nathalie Tzourizio-Mazoyer, Bernard Tzourizio-Mazoyer and Sophie Maingault for their generosity in sharing the surface template created in the BIL&GIN; a unique database enriched in left-handed individuals for the study of human brain lateralization.

## Disclosure statement

The authors have no conflicts of interest to disclose.

## Notes

### Competing Interest Statement

The authors have declared no competing interest.

