## Supplementary material for "Asymmetric thinning of the cerebral cortex across the adult lifespan is accelerated in Alzheimer’s Disease": SI Fig 1

**Supplementary information**

1. Methods

### 1. Methods

#### *1.1 Samples*

The main discovery sample consisted of magnetic resonance imaging (MRI) data collected across 5 projects at the Center for Lifespan Changes in Brain and Cognition (LCBC; Department of Psychology, University of Oslo): Neurocognitive Development [1]; Cognition and Plasticity Through the Lifespan [2]; Constructive Memory [3]; Method of Loci [4]; and Neurocognitive Plasticity [5]. Because the latter two studies involved cognitive training, only MRI data from these projects was considered. The details of the individual studies have been

| Sample | N unique | N obs | N Longitudinal | Mean Time Interval (SD) | Interval Range | N Timepoints | Mean Age (range) | Sex (F/M) |
|---|---|---|---|---|---|---|---|---|
| LCBC (discovery) | 1084 | 2577 | 1851 | 2.7(2.8) | 0.1 - 11 | 1-6 | 50.0(20.0-89.4) | 703/381 |
| Cam-CAN | 634 | 898 | 528 | 1.4(0.7) | 0.2 - 3.5 | 1-2 | 55.5(20.2-91.6) | 323/311 |
| BASE-II | 447 | 768 | 642 | 1.9(0.7) | 0.6 - 3.1 | 1-2 | 62.4(24.1-83.1) | 170/277 |
| BETULA | 310 | 480 | 340 | 4(0.2) | 3.4 - 4.6 | 1-2 | 62.7(25.5-84.8) | 159/151 |
| DLBS | 471 | 763 | 584 | 3.9(0.4) | 2.5 - 5.6 | 1-2 | 59.7(20.6-93.1) | 292/179 |

described at length elsewhere (see [6]). For 563 individuals, more than one scan was available; 1851 of these were longitudinal in nature (mean number of longitudinal scans per participant = 3.3), whereas 205 were double-scanned at the same timepoint on both LCBC scanners (1.5T Avanto and 3T Skyra; Siemens Medical Solutions). The number of timepoints ranged from 1 to 6 (mean = 3.3 ± 1.8) and the max follow-up interval was 11.0 years since initial scan. For cognitive analyses, cognitive testing was conducted on average 19.0 days post-scan and all were tested within 5 months.

*SI Table 1: Description of the discovery sample (LCBC) and four replication samples. Intervals are in years.*

| | | | N Timepoints | | | | | | | | Mean Clinical Dementia Rating |
|---|---|---|---|---|---|---|---|---|---|---|---|
| **Group** | **N unique** | **N Obs** | **2** | **3** | **4** | **Mean Time Interval (SD)** | **Interval Range** | **Mean Age (range)** | **Sex (f/m)** | **All timepoints** | **Longitudinal timepoints** |
| NC-long | 128 | 435 | 22 | 33 | 73 | 2.1 (1.4) | 1.05 - 6.8 | 73.0 (60.5 - 90.3) | 221/214 | 0.02 | 0.01 |
| AD-long | 41 | 110 | 20 | 14 | 7 | 1.6 (1.1) | 1.35 - 5.4 | 74.7 (55.0 - 89.1) | 55/55 | 0.8 | 0.93 |

*SI Table 2: Description of the AIBL AD sample groups. Intervals are in years.*

| Sample | Scanner | Sequence | Tesla | Head coil | Slices | Voxel size (mm) | Time parameters | Other parameters |
|---|---|---|---|---|---|---|---|---|
| LCBC (discovery) | Avanto (Siemens) | 3D MP-RAGE | 1.5 | 12-channel | 160 | 1.25×1.25×1.25 | TR/TE/TI=2400ms/3.61ms/1000ms | FA/FOV = 8°/240×240m |
| | Skyra (Siemens) | 3D MP-RAGE | 3 | 32-channel | 176 | 1×1×1 | TR/TE/TI=2300ms/2.98ms/850ms | FA/FOV = 8°/256×256m |
| Cam-CAN | Tim Trio (Siemens) | 3D MP-RAGE | 3 | 32-channel | 192 | 1×1×1 | TR/TE/TI=2250ms/2.98ms/900ms | FA/FOV = 9°/256×240m |
| BASE-II | Tim Trio (Siemens) | 3D MP-RAGE | 3 | 32-channel | 176 | 1×1×1 | TR/TE/TI=2500ms/4.77ms/1100ms | FA/FOV = 7°/256×256m |
| BETULA | Discovery (GE) | 3D FSPGR | 3 | 32-channel | 176 | 1×1×1 | TR/TE/TI=8.19ms/3.2ms/450ms | FA/FOV = 12°/250×250r |
| DLBS | Achieva (Philips) | 3D MP-RAGE | 3 | 8-channel | 160 | 1×1×1 | TR/TE/TI=2300ms/8.13ms/1100ms | FA/FOV = 12°/204×256r |
| AIBL (AD sample) | Avanto (Siemens) | 3D MPRAGE | 1.5 | **NA** | 160 | 1×1×1.2 | TR/TE/TI=2300ms/2.98ms/900ms | FA/FOV = 9°/240×256m |
| | Verio (Siemens) | 3D MPRAGE | 3 | **NA** | 160 | 1×1×1.2 | TR/TE/TI=2300ms/2.98ms/900ms | FA/FOV = 9°/240×256m |
| | TrioTim (Siemens) | 3D MPRAGE | 3 | **NA** | 160 | 1×1×1.2 | TR/TE/TI=2300ms/2.98ms/900ms | FA/FOV = 9°/240×256m |

*SI Table 3 MRI acquisition parameters by sample. TR =Repetition time; TE= Echo time; TI = Inversion time; FA = Flip angle; F'OV = Field of view; NAV = not available [7]; 3D MPRAGE = three-dimensional magnetization prepared rapid gradient echo; 3D FSPGR = three-dimensional fast spoiled gradient echo.*

##### *1.2 MRI preprocessing*

First, T1w data was processed in FreeSurfer 6.0 cross-sectionally, which includes steps such as removal of non-brain tissue, Talairach transformation, intensity normalisation, demarcating the grey/white and grey/CSF boundaries, and reconstructing the cortical surface. This automated pipeline yields a reconstructed surface map and cortical thickness estimates for each person at each timepoint. To extract more reliable thickness estimates, the reconstructed images were processed longitudinally. Specifically an unbiased within-subject template space and image is created using robust, inverse consistent registration [8] Several processing steps, such as skull stripping, Talairach transforms, atlas registration as well as spherical surface maps and parcellations are then initialized with common information from the within-subject template, significantly increasing reliability and statistical power of the cortical thickness estimates [9].

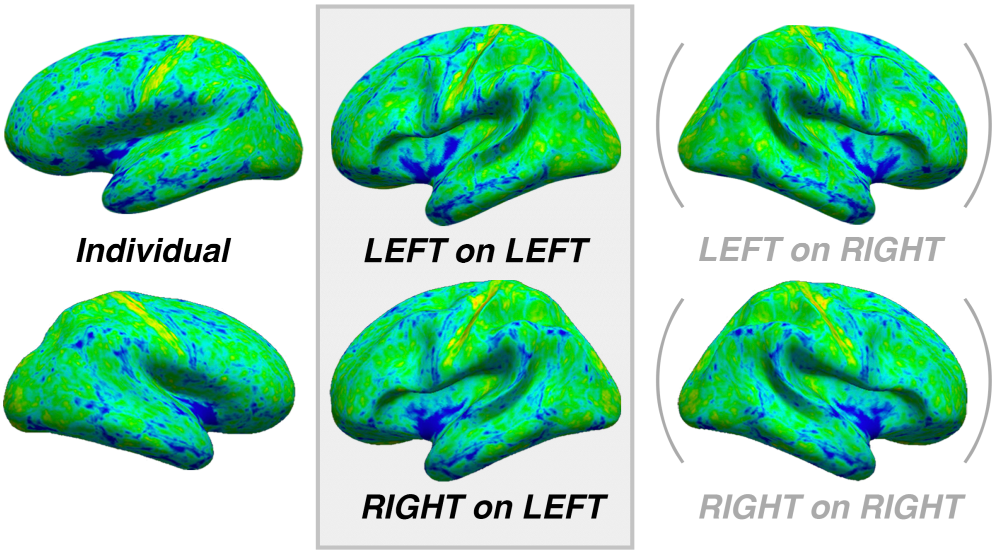

*SI Fig. 1. Left and right cortical thickness maps of an example subject shown on the individual surface (see left), and after resampling the data from both hemispheres into homotopic alignment using either one hemisphere of the symmetrical surface. All analyses were performed after resampling to the left symmetrical surface (middle). For visual comparison data is also shown on the right symmetrical surface (right).*

#### 2. Results

##### *2.1 Comparison with vertex-wise Linear Mixed Effects (discovery sample)*

Conventional vertex-wise linear mixed effect (LME) analyses assessing the linear Age × Hemisphere interaction (F-test with Age, $Age^2$,$Age^3$) yielded near-identical results to the Age × Hemisphere GAMM interaction (SI Fig.2A; Dice coefficient = 0.84; 96% of significant vertices were encompassed within LME effects), indicating significance results from our non-linear approach were not due to overfitting.

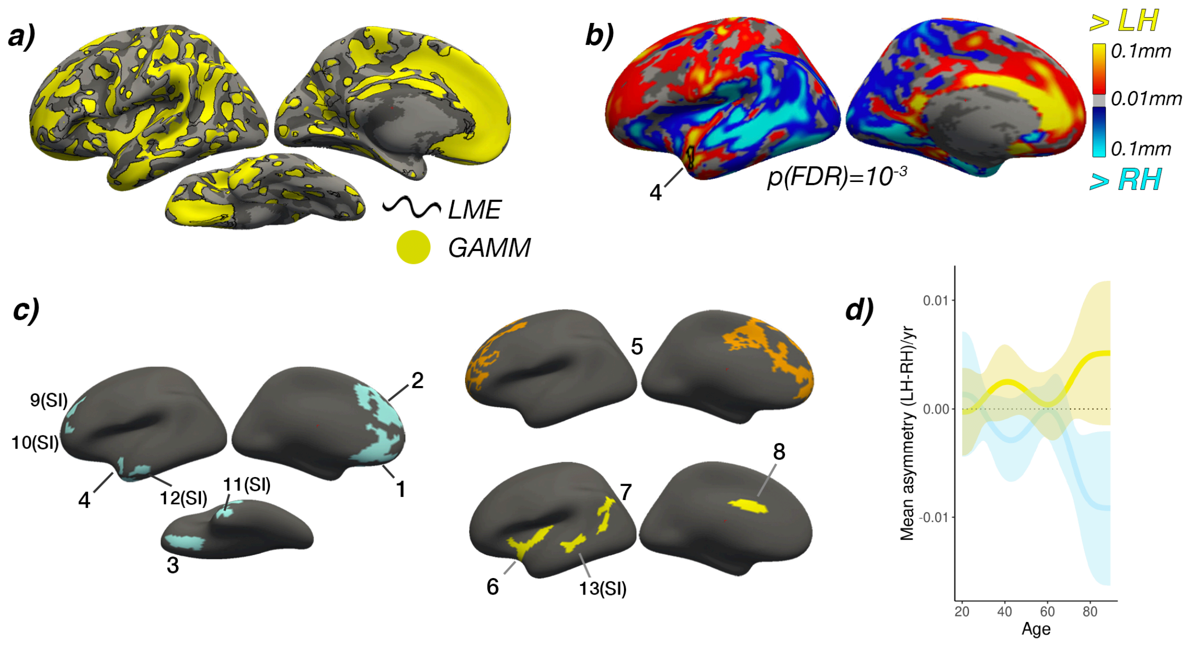

*SI Fig.2): **a)** comparison of significance maps between our non-linear Generalized Additive Mixed Models (GAMM) approach (yellow; age-changes in asymmetry GAMM effects and a conventional linear mixed model (LME) approach (black outline). 96% of vertices exhibiting significant GAMM effects were encompassed within vertices showing significant LME effects. **B)** Hemisphere effect in the main sample (visualized in main paper) after resampling to fsaverage5 to permit visual comparison with clustering results. ROI #4 is shown to illustrate the overlap with asymmetry effects in the temporal lobe. **C)** The ROI's derived from clustering (numbered 1-13). The trajectories of ROI's marked with SI are not shown in the main paper but can be seen in SI Fig.4). **D)** The first derivative of the asymmetry trajectories for cluster solutions 1 and 3 in the LCBC discovery sample. The mean timeline of change was highly similar for ROI's exhibiting LH-asymmetry loss and RH-asymmetry loss across the adult lifespan.*

#### *2.2 Clustering of asymmetry trajectories (discovery sample)*

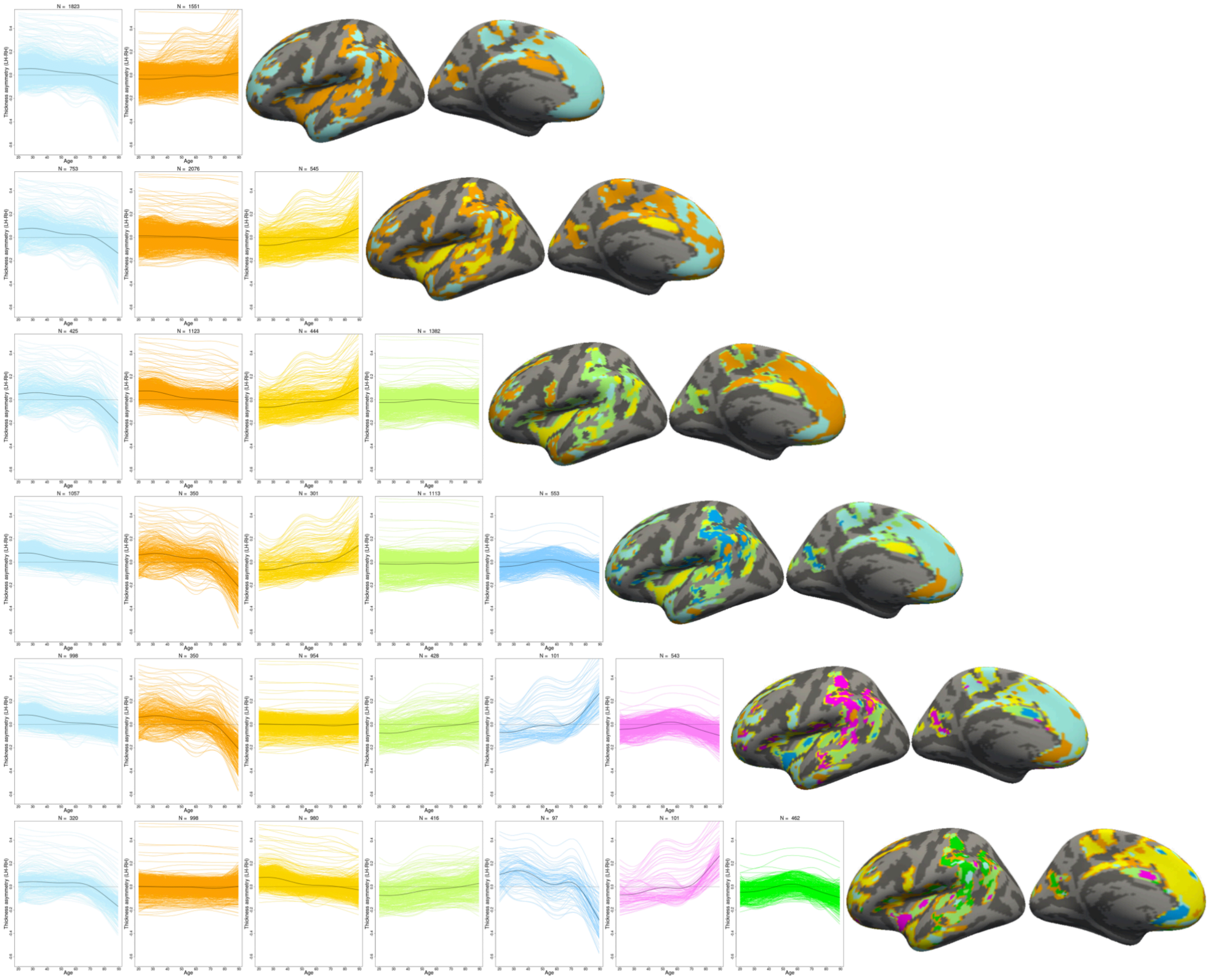

*SI Fig.3: The main findings were robust to varying the number of PAM clustering partitions (LCBC sample).*

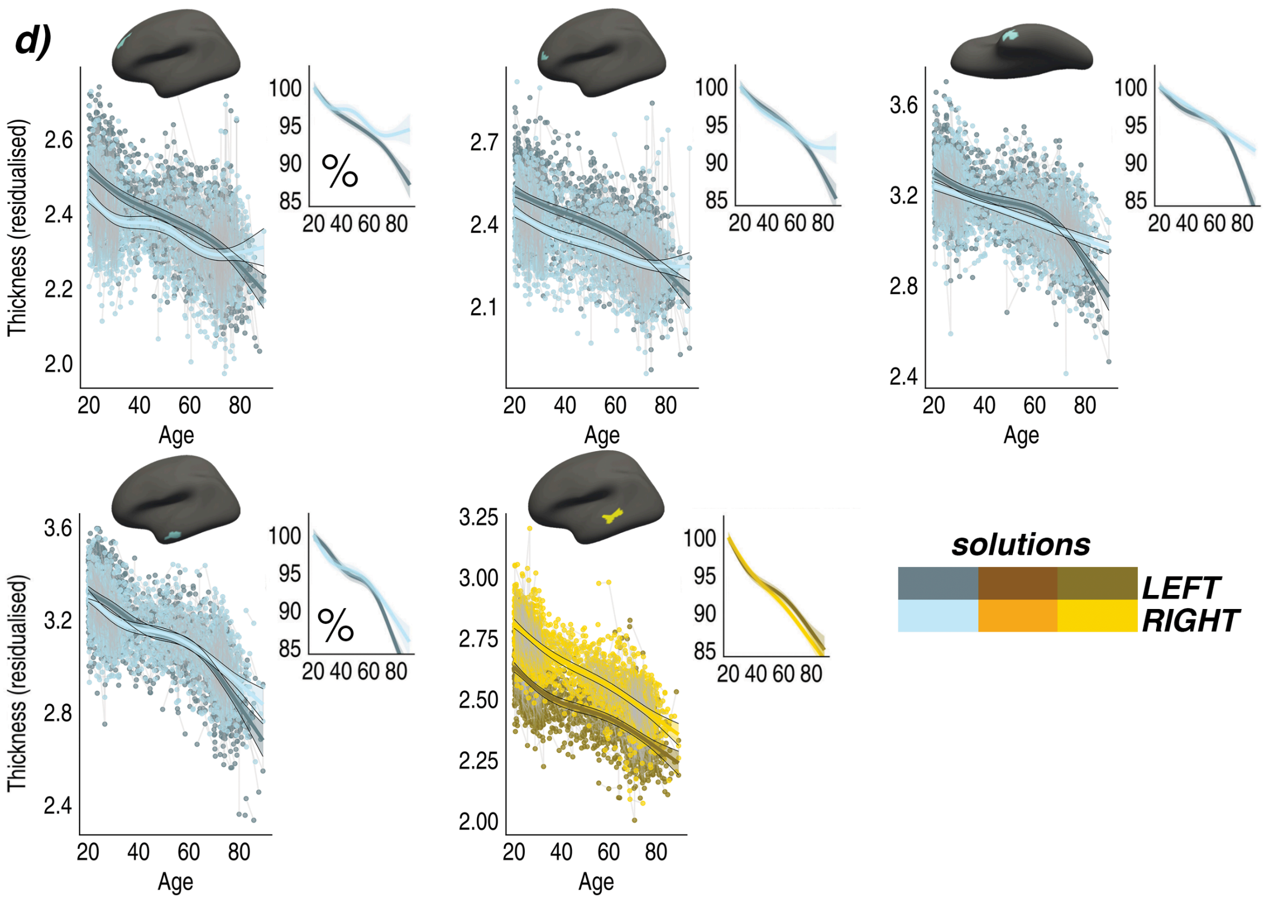

*SI Fig.4. Age-trajectories of cortical thickness plotted separately for the left and right hemisphere for the remaining clustering-derived regions. All trajectories were fitted using Generalized Additive Mixed Models (GAMMs). Colors correspond to the solutions in Fig.3A (main paper), and darker shades indicate left hemisphere trajectories. Data is residualized for sex and scanner. Ribbons depict 95% confidence intervals. Smaller plots illustrate percentage-change with age for each region.*

##### *2.3 Comparison of asymmetry trajectories between discovery and replication samples*

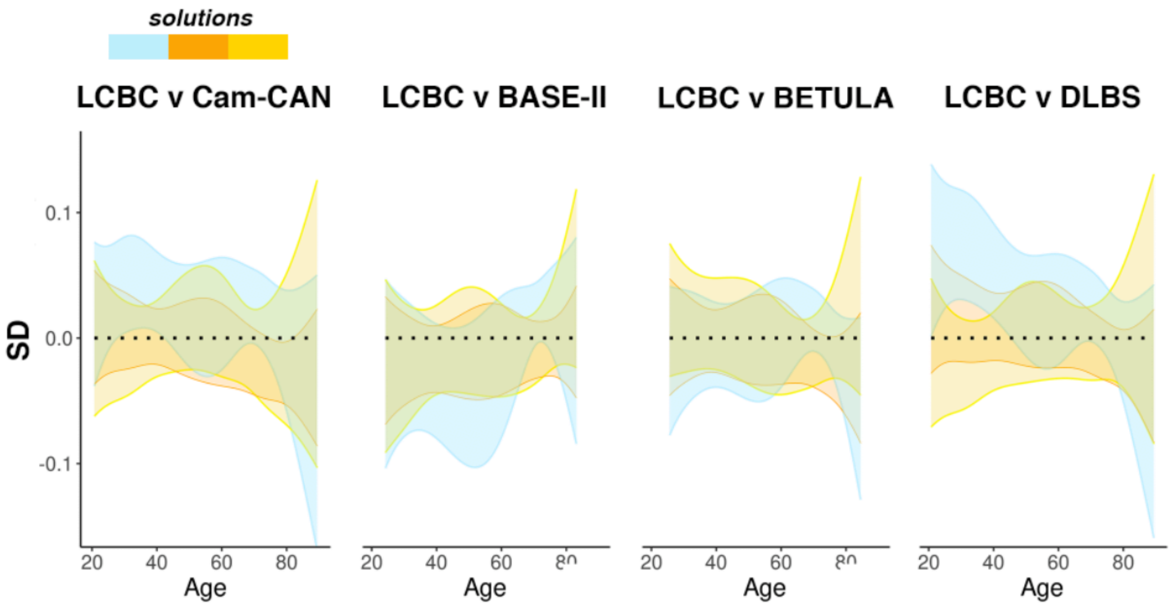

*SI Fig.5. Comparison of solutions for mean asymmetry trajectories between replication and discovery samples. The means and standard deviations of the three clustering solutions in each sample were subtracted from those in the LCBC discovery sample. Standard deviation was used as the measure is not influenced by the (arbitrary) number of vertices on the surface.*

##### *2.4 Vertex-wise analysis in replication samples*

Vertex-wise GAMM Age × Hemisphere effects appeared largely consistent in Cam-CAN, BASE-II and to a lesser extent BETULA (SI Fig.6). Spatial correlation of effect size maps with LCBC confirmed the impression that vertex-wise significance of asymmetry-change effects replicated in Cam-CAN and BASE-II (respectively r = .45, r = .52), though did not substantiate a replication of vertex-wise effects in BETULA (r = .01) or DLBS (r = .08). The main effect of Hemisphere was highly reproducible across samples (SI Fig. 7).

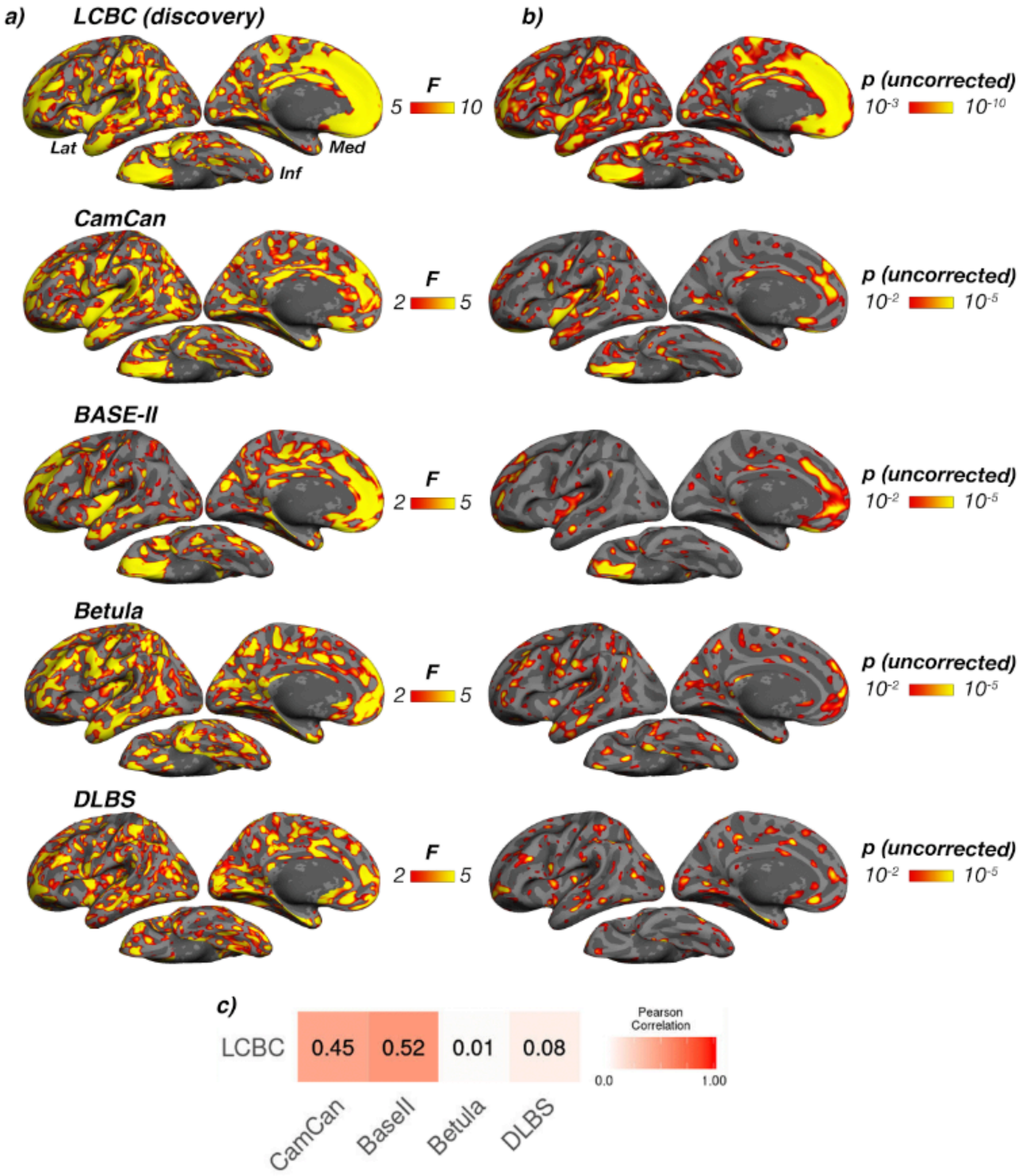

*SI Fig.6. Vertex-wise GAMM age-change in asymmetry effects for all samples. Samples are depicted row-wise. Note that for visualization purposes, a slightly different threshold is shown for LCBC data to account for differences in sample size/test power.* ***A)*** *F values* ***B)*** *significance effects (uncorrected).* ***C)*** *spatial correlation of effect size maps ($\Omega^2$; maps not shown) for each sample against $\Omega^2$ in the main LCBC discovery sample. $\Omega^2$ was used to assess overlap as the relative values are unaffected by differences in sample size/test power. Spatial correlation between discovery and replication samples was quantified using Pearson's r. Lat = lateral; med = medial; inf = inferior.*

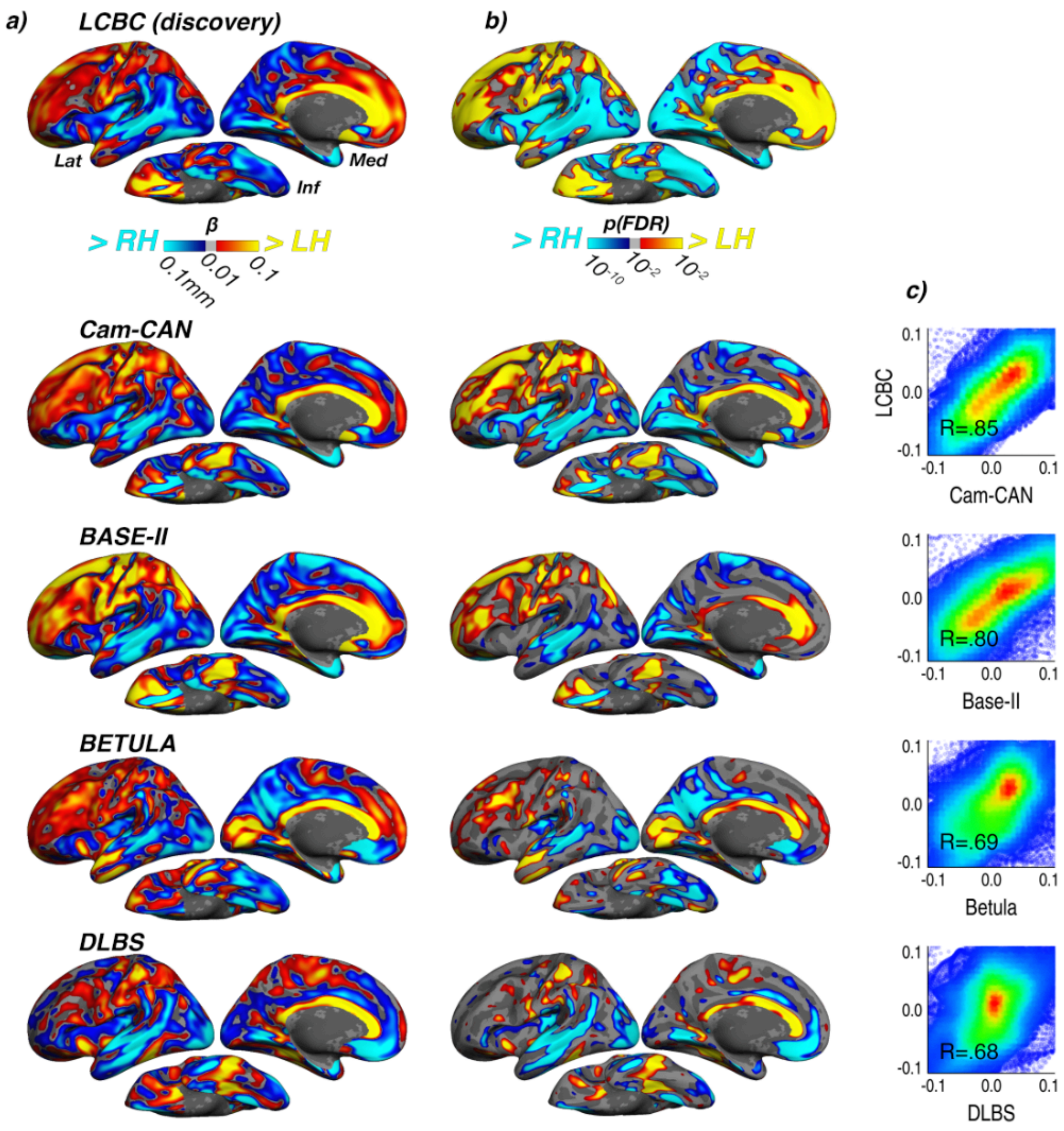

*SI Fig.7. Vertex-wise main effect of Hemisphere for all samples. Samples are depicted row-wise.* ***A)*** *Beta coefficients in mm* ***B)*** *significance effects (FDR corrected). Note that differences in sample size (and consequently test power) between samples affect the overall level of significance in column B but not A.* ***C)*** *spatial correlation of Beta coefficients for each sample against Beta coefficients in the main LCBC discovery sample. Spatial correlation between discovery and replication samples was quantified using Pearson's r. Lat = lateral; med = medial; inf = inferior.*

##### *2.5 Clustering analysis in replication samples*

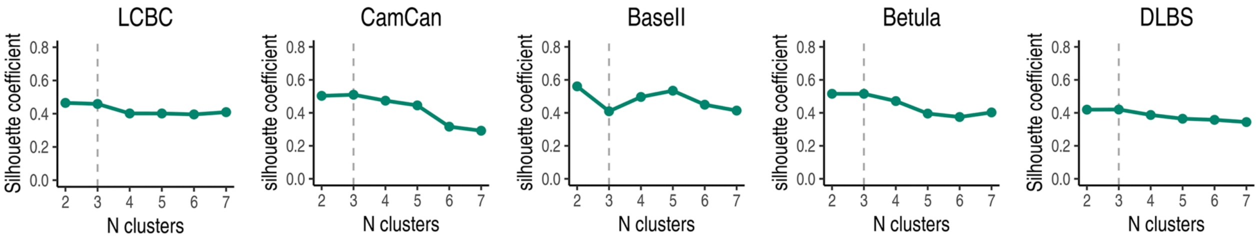

*SI Fig.8. Mean silhouette width used to determine the optimal number of clustering partitions, here shown plotted for each cohort. The chosen 3-cluster solution (identified in the LCBC discovery sample and subsequently applied in replication cohorts) showed stability across samples. Note that BASE-II is the only cohort with non-continuous age sampling which will affect the estimation and subsequent clustering of trajectories (cf. Fig. 1 in main paper).*

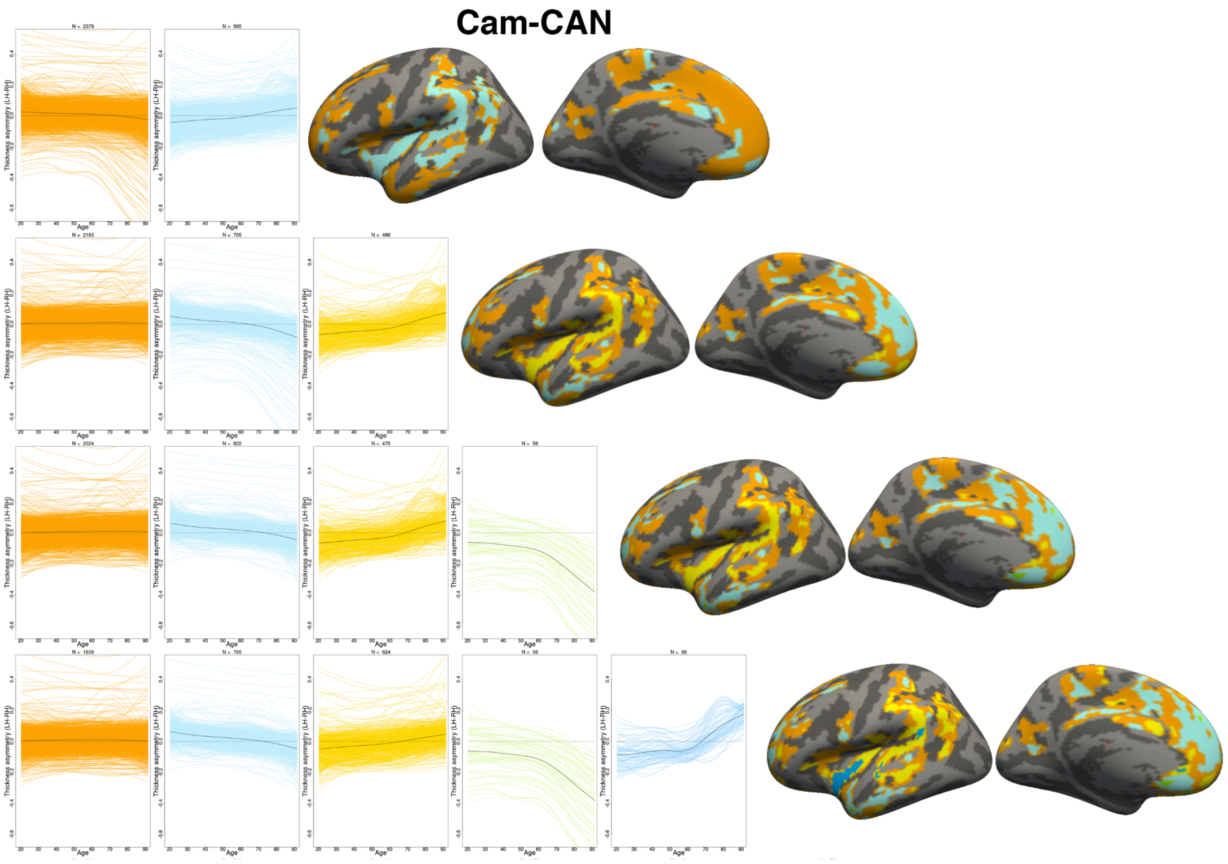

*SI Fig.9 (above and continued below): The main findings were conserved when varying the number of PAM clustering solutions also in replicating cohorts. Note that colors of the clustering maps are not comparable across cohorts because the order in which clusters are identified by the algorithm will necessarily differ across cohorts.*

*SI Fig.9 (continued):*

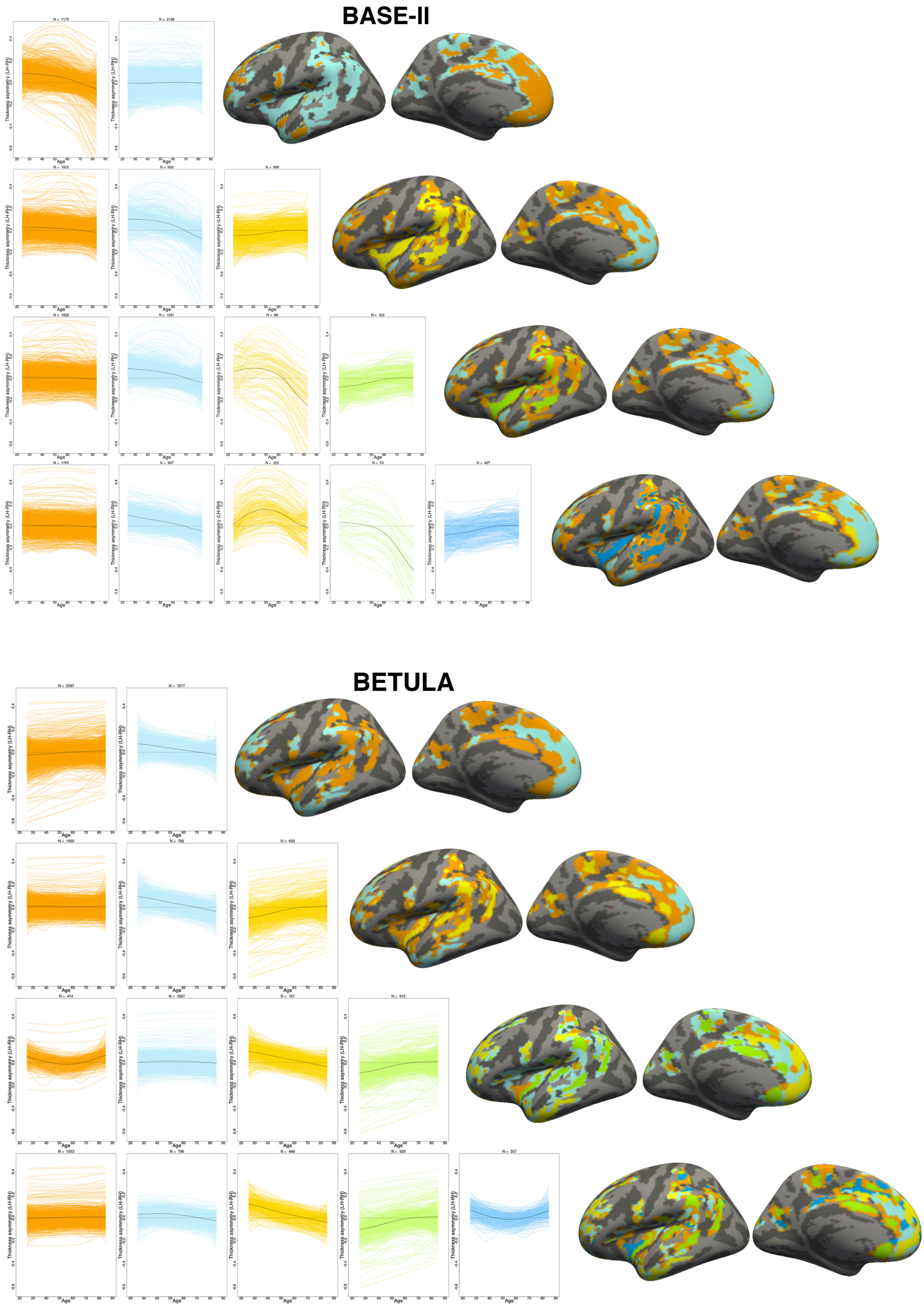

*SI Fig.9 (continued):*

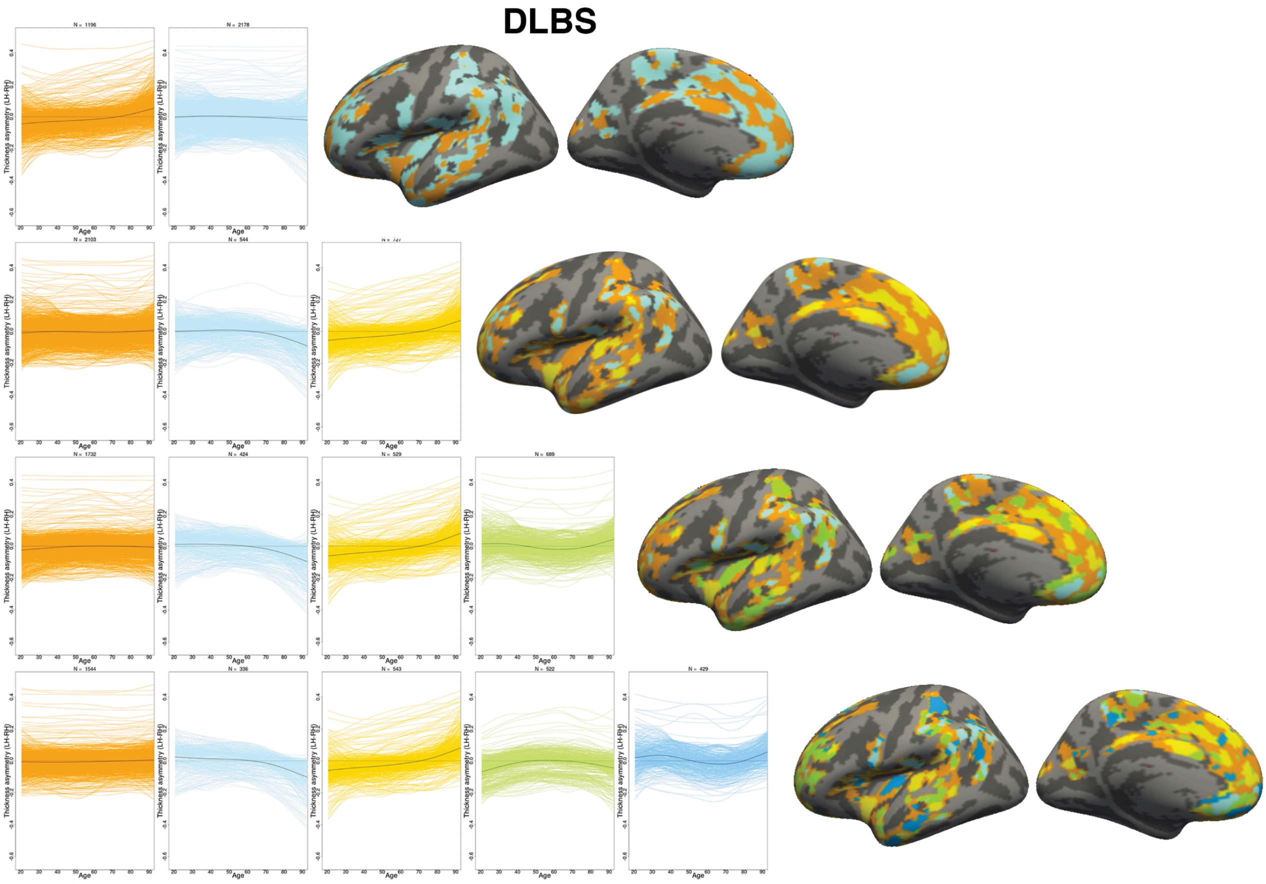

##### *2.6 Cognitive change analysis*

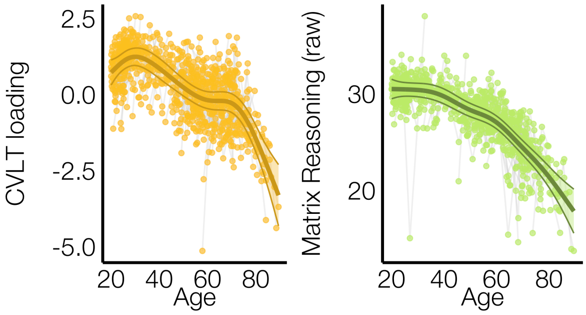

*SI Fig. 10. GAMMs revealed that Age (Sex corrected) explained 29% and 36% of the variance in changes in memory (CVLT) and fluid reasoning ability (Matrix reasoning), respectively. Data is residualized for sex and (CVLT) test version. See SI Table 2 for full stats.*

| | Age effect | | | Covariate | Model | |
|---|---|---|---|---|---|---|
| | *edf* | *F* | *p-uncorrected* | *Sex (p-uncorrected)* | $r^{2\ (adjusted)}$ | $\Omega^2$ |
| CVLT | 6.12 | 25.57 | $10^{-28}$ | <0.001 | 0.29 | 0.16 |
| Matrix reasoning | 4.61 | 58.45 | $10^{-49}$ | 0.5 | 0.36 | 0.25 |

*SI Table 3. Full GAMM results predicting changes in verbal memory (CVLT) and fluid reasoning ability (Matrix reasoning) as a function of Age (Sex-corrected)*

.

| **ROI #** | **Region** | **Clustering solution** | **Cognitive variable** | **Thickness-asymmetry (LH-RH)** | | | | **Covariates** | | | **Model without Age covariate** | |
|---|---|---|---|---|---|---|---|---|---|---|---|---|
| | | | | *edf* | *F* | *p-uncorrected (Effect direction)* | *p(FDR)* | *Mean thickness (p-uncorrected [effect direction])* | *Sex (p-uncorrected)* | *Scanner (p-uncorrected)* | *Thickness-asymmetry (LH-RH) p-uncorrected (Effect direction)* | *Mean thickness (p-uncorrected [effect direction])* |
| 1 | rostral anterior cingulate | 1 | CVLT | 3.34 | 1.33 | 0.37 | 0.72 | 0.66 | <0.001 | <0.01 | 0.6 | <0.01 (thicker) |
| 2 | superior frontal | | | 2.14 | 2.11 | 0.1 | 0.39 | 0.36 | <0.001 | <0.001 | 0.69 | <0.001 (thicker) |
| 3 | lateral orbitofrontal | | | 1 | 5.29 | 0.02 (lower) | 0.29 | 0.81 | <0.01 | <0.001 | 0.33 | <0.001 (thicker) |
| 4 | superior temporal | | | 1 | 0.37 | 0.54 | 0.72 | 0.04 (thinner) | <0.001 | <0.01 | 0.05 | 0.001 (thicker) |
| 5 | frontal cortex | 2 | | 1 | 4.39 | 0.04 (lower) | 0.29 | 0.65 | <0.001 | <0.01 | 0.97 | <0.001 (thicker) |
| 6 | insula | 3 | | 1 | 0.33 | 0.57 | 0.72 | 0.53 | <0.001 | <0.01 | 0.01 (higher) | 0.001 (thicker) |
| 7 | inferior parietal | | | 1 | 0.5 | 0.48 | 0.72 | 0.86 | <0.001 | <0.01 | 0.36 | <0.001 (thicker) |
| 8 | caudal anterior cingulate | | | 1 | 0.18 | 0.67 | 0.73 | 0.77 | <0.001 | <0.01 | 0.06 | 0.496 |
| 1 | rostral anterior cingulate | 1 | Matrix Reasoning | 1 | 0.3 | 0.59 | 0.72 | 0.09 | 0.6 | 0.66 | 0.03 (higher) | <0.001 (thicker) |
| 2 | superior frontal | | | 1.54 | 0.25 | 0.69 | 0.73 | 0.03 (thicker) | 0.37 | 0.33 | 0.07 | <0.001 (thicker) |
| 3 | lateral orbitofrontal | | | 1 | 0.63 | 0.43 | 0.72 | 0.08 | 0.54 | 0.54 | <0.001 (higher) | <0.001 (thicker) |
| 4 | superior temporal | | | 1.92 | 3.5 | 0.06 | 0.34 | 0.64 | 0.63 | 0.3 | <0.001 (higher) | <0.001 (thicker) |
| 5 | frontal cortex | 2 | | 1 | 1.55 | 0.21 | 0.65 | 0.07 | 0.43 | 0.44 | 0.14 | <0.001 (thicker) |
| 6 | insula | 3 | | 2.23 | 0.88 | 0.32 | 0.72 | 0.05 | 0.56 | 0.25 | 0.001 (higher) | <0.001 (thicker) |
| 7 | inferior parietal | | | 1 | 0.04 | 0.85 | 0.85 | 0.85 | 0.52 | 0.5 | 0.27 | <0.001 (thicker) |
| 8 | caudal anterior cingulate | | | 2.12 | 1.4 | 0.25 | 0.65 | 0.03 (thicker) | 0.31 | 0.92 | 0.16 | 0.021 (thicker) |

*SI Table 4: Longitudinal cognitive analyses predicting age-changes in verbal memory (California Verbal Learning Test [CVLT]) and fluid reasoning (Matrix Reasoning subtest of WAIS). GAMM analyses were conducted assessing the predictive value of the smooth term (s) of s(Thickness asymmetry;[LH-RH]) controlling for s(Age), s(Mean thickness; [LH+RH/2]), Sex and Scanner. No significant effects of Thickness asymmetry were observed after FDR-correction (16 comparisons) in any of the 8 ROI's for either cognitive measure. Estimated* *degrees of freedom (edf) is an index of curve complexity (i.e. edf = 1 approximates a linear association). P-values pre- and post FDR-correction are shown for Thickness asymmetry effects (columns 7-8) . The direction of the asymmetry effect is given for uncorrected p-values < .05 (e.g. higher = more asymmetry relates to better cognition; lower = less asymmetry relates to better cognition). Effect directions for Mean thickness are also given (thicker = thicker cortex relates to better cognition; thinner = thinner cortex relates to better cognition). Covariate p-values are shown in right columns. Results of comparable GAMM models without covarying for Age are shown in the rightmost columns. Significant asymmetry effects were observed when covarying for mean thickness (across hemispheres) but not age. Uncorrected p-values and directions for the effects of s(Thickness Asymmetry) and Mean thickness are reported. The total number of observations for CVLT and Matrix Reasoning with concurrent scans were 783 (N = 312) and 788 (N = 314), respectively.*

Having observed no significant effects of thickness asymmetry change upon longitudinal cognitive change in any of the 8 ROI's alone, we finally assessed whether regional changes in thickness asymmetry nevertheless contributed a significant combined effect upon cognitive change. Here, we assessed the difference in model fit between a GAMM containing the smooth terms of thickness asymmetry for all 8 ROI's versus one without (covaried for Age, mean thickness [here the first two principal components of the mean thickness of all 8 ROI's], sex and scanner). No significant difference in model fit was observed (verbal memory p = .98; fluid reasoning p = .79), indicating asymmetry had no significant combined effect upon cognitive change.

##### *2 .7 Longitudinal Alzheimer's disease (AD) analysis*

| ROI # | Region | Clustering solution | df | NC-long v AD-long | | | |
|---|---|---|---|---|---|---|---|
| | | | | *T* | *p-uncorrected* | *p(FDR)* | *Sig* |
| 1 | rostral anterior cingulate | 1 | 374 | -2.44 | 0.02 | 0.04 | * |
| 2 | superior frontal | | 374 | -1.67 | 0.095 | 0.152 | |
| 3 | lateral orbitofrontal | | 374 | -4.65 | <0.001 | <0.001 | * |
| 4 | anterior temporal | | 374 | -3.16 | 0.002 | 0.007 | * |
| 5 | frontal cortex | 2 | 374 | -2.3 | 0.022 | 0.044 | * |
| 6 | insula | 3 | 374 | -1.06 | 0.289 | 0.385 | |
| 7 | inferior parietal | | 374 | 0.83 | 0.41 | 0.468 | |
| 8 | caudal anterior cingulate | | 374 | 0.23 | 0.817 | 0.817 | |

*SI Table 5: Longitudinal AD analysis. Change-change analyses were conducted assessing the Group x Years (since baseline) interaction, indicating group differences in thickness-asymmetry change over time between AD- and NC-classified individuals. Significant FDR-corrected effects (i.e. p(FDR) < 0.05) were observed in frontal cortical and anterior temporal regions. P-values pre- and post FDR correction are shown. AD = Alzheimer's diease; NC = Normal controls; Sig = significance.*

#### *4. Supplementary R Code*

```
require("dplyr")
require("gamm4")

#vertex-wise
for (j in 1:end) {
  if (j == 1) {

    # make 100 obs age model
    nn=100
    Opt <- list()
    Opt$Age <- seq(min(age), max(age), length.out = 100)
    Opt$fake.frame = data.frame("Age" = Opt$Age,
                                "hemi" = rep(1,nn),
                                "sex_demean" = rep(0,nn),
                                "scanner_demean" = rep(0,nn))
    pdat = rbind(Opt$fake.frame,Opt$fake.frame)
    pdat$hemi[1:100] = 0

    #set outputs
    gamm.trajectories <- list()
    ogamm.trajectories <- list()
    RR <- list()
    RR$edf <- NULL
    RR$Fval <- NULL
    RR$p.spl <- NULL
    RR$pp.spl <- NULL
    RR$fit_valL <- NULL
    RR$fit_valR <- NULL
    RR$fit_valdiff <- NULL
    RR$se <- NULL
    RR$hT <- NULL
    RR$hCoef <- NULL
    RR$hP <- NULL
    RR$hPlog <- NULL
  }

  gamm.trajectories[[j]] = gamm4(thickness ~ s(Age, by = as.factor(hemi), k = 6) +
            as.factor(hemi) + sex_demean + scanner_demean,
            data = dat,
            random = ~ (1 |subID))
  gamm.sum = summary(gamm.trajectories[[j]]$gam)
  g = gamm.trajectories[[j]]$gam

  #simulate from posterior - Xp matrix
  #muliply matrix by model coefficients and sum rows to get predicted values
  Xp = predict(g, newdata = pdat, type = "lpmatrix")
  Xp %*% coef(g)
```

```
#which cols of Xp relate to which hemi
c1 = grepl("hemi\\)1", colnames(Xp)) #L
c2 = grepl("hemi\\)0", colnames(Xp)) #R

#which rows of Xp
r1 = with(pdat, hemi == 1) #L
r2 = with(pdat, hemi == 0) #R

#differences between smooths
#cols of differenced Xp matrix that aren't involved in comparison set to zero
X <- Xp[r1, ] - Xp[r2, ] #L-R
X[, !(c1 | c2)] <- 0
X[, !grepl("s\\(", colnames(Xp))] = 0

#get predicted vals of zero-centred hemispheric difference at zero of other covs
#SE of the difference using variance-covariance matrix of
#estimated model coefficients
dif <- X %*% coef(g)
se <- sqrt(rowSums((X %*% vcov(g, unconditional =T)) * X))

#ordered factor approach to get test statistics for GAMM interaction
dat = mutate(dat,
  ohemi = ifelse(dat$hemi == 1, "left", "right"),
  ohemi = factor(ohemi, levels = c("left","right"),ordered = T))

#estimate smooth for set reflevel and
#smoothed difference between ref and other levels
ogamm.trajectories[[j]] = gamm4(thickness ~ as.factor(hemi) +
          s(Age) + s(Age, by = ohemi, k = knots) +
          sex_demean + scanner_demean, data = dat,
          random = ~ (1 |subID))
ogamm.sum = summary(ogamm.trajectories[[j]]$gam)

#get stats
RR$edf = RR$edf %>% cbind(.,
                      ogamm.sum$edf[2])
RR$Fval = RR$Fval %>% cbind(.,
                      ogamm.sum$s.table[[2,3]])
RR$p.spl = RR$p.spl %>% cbind(.,
                      -log10(ogamm.sum$s.pv[2]))

#get fits
RR$fit_valL = RR$fit_valL  %>%  cbind(.,
          predict.gam(gamm.trajectories[[j]]$gam, newdata = pdat[101:200,])) #L
RR$fit_valR = RR$fit_valR  %>%  cbind(.,
```

```
            predict.gam(gamm.trajectories[[j]]$gam, newdata = pdat[1:100,])) #R
  RR$fit_valdiff = RR$fit_valdiff  %>%  cbind(., dif)
  RR$se = RR$se %>% cbind(., se)

  #hemieffects
  RR$hT = RR$hT %>% cbind(., gamm.sum$p.t[[2]])
  RR$hCoef = RR$hCoef %>% cbind(., gamm.sum$p.coeff[[2]])
  RR$hP = RR$hP %>% cbind(., gamm.sum$p.pv[[2]])
  RR$hPlog = RR$hPlog %>% cbind(., -log10(gamm.sum$p.pv[[2]]))

}

#for further reading see
#https://fromthebottomoftheheap.net/2017/10/10/difference-splines-i/
#https://fromthebottomoftheheap.net/2017/12/14/difference-splines-ii/
```
